# 2-hydroxyglutarate controls centromere and heterochromatin conformation and function in the male germline

**DOI:** 10.1101/2022.05.29.493890

**Authors:** Nina Mayorek, Miriam Schlossberg, Yousef Mansour, Nir Pillar, Ilan Stein, Fatima Mushasha, Guy Baziza, Eleonora Medvedev, Zakhariya Manevitch, Julia Menzel, Elina Aizenshtein, Boris Sarvin, Nikita Sarvin, Tomer Shlomi, Michael Klutstein, Eli Pikarsky

## Abstract

2-hydroxyglutarate (2HG) is recognized as an epigenetic regulator in cancer and some transient biological processes. Of all organs, the testis harbors the highest baseline physiological levels of 2HG, yet it’s putative functions in germ cell biology are unknown. Here we show that 2HG is generated in specific stages of spermatogenesis by the testis specific lactate dehydrogenase C (LDHC), beginning at the last stages of prophase I. Unexpectedly LDHC enters nuclei and concentrates in centromeres. LDHC-generated L-2HG controls centromere condensation and pericentromeric heterochromatin organization through multiple effects including clustering of chromocenters, centromere and chromocenter condensation and expression of satellite RNAs. These effects are rapid and specific to L but not D-2HG. *In vivo* depletion of L-2HG causes centromere malfunction and activation of the spindle assembly checkpoint. Our findings reveal that 2HG can directly affect centromere and pericentromeric heterochromatin conformation and function and is necessary for licensing chromosome segregation.

## Introduction

Germ cell differentiation in the testis involves major phenotypic, transcriptional, regulatory, metabolic and epigenetic changes. These changes are programmed to allow the transmission of genetic information to the next generation through highly complex, multistage differentiation processes beginning from the diploid spermatogonial-stem-cell stage to haploid and highly specialized spermatozoa. Few studies have directly addressed the role of metabolic changes in germ cells during mammalian germ cell differentiation and meiosis^1,2^. One well-studied example is the effect of retinoic acid (RA) and its metabolism on meiosis entry^3,4^. Changes in the metabolic state of the cell can affect its epigenetic landscape, dictate cell function and may have heritable consequences^5,6^. It is thus conceivable that cellular metabolism may have far-reaching effects on meiosis progression and germ cell differentiation.

2-hydroxyglutarate (2HG) arises from the reduction of α-ketoglutarate (α-KG) in several pathophysiological systems^7^. There are two enantiomers: D-2HG acquired by the neomorphic catalytic activity of mutated isocitrate dehydrogenase IDH1/2^8^ and L-2HG which is derived from promiscuous activity of a number of enzymes, including lactate dehydrogenase A (LDHA)^9^, the testis specific LDHC^10^, and malate dehydrogenase (MDH)^11^. These metabolites are degraded by the activity of the enantiomer specific 2HG dehydrogenases and mutations in these enzymes cause an accumulation of 2HG resulting in metabolic diseases.

Accumulation of 2HG in certain tumors and its involvement in tumorigenesis identified both enantiomers as oncometabolites. Gain of function mutations in IDH1/2 cause tumors in multiple organs, including glioma, leukemia, cholangiocarcinoma and chondrosarcoma via formation of high concentrations of D-2HG^12,13^. Similarly, in clear-cell renal-cell-carcinoma copy number loss of L-2HGDH together with promiscuous activity of MDH leads to the accumulation of L-2HG^14^. Both enantiomers of 2HG are inhibitors of α-KG/Fe (II)-dependent dioxygenases, including the JmjC-domain family of histone demethylases and the TET family of DNA demethylases. Thus, it was deduced that 2HG is likely ‘oncogenic’ via the dysregulation of the epigenetic state of cell. In addition to these pathological conditions, L-2HG was found to play a role in cell fate decisions of CD8^+^ T lymphocytes in response to T cell receptor triggering. In these cells, global DNA methylation and changes in certain histone methylations caused changes in gene expression^17^. L-2HG also plays a role in the adaptation to hypoxia^9,18^. In this case, intracellular acidification resulting from limited oxygen supply extends substrate specificity of LDHA and MDH allowing these enzymes to use αKG as a substrate for L-2HG formation. This helps in alleviating hypoxia-induced reductive stress by using NADH to convert α-KG into L-2HG^18^.

The testis is unique in accumulating 2HG in non-pathological conditions, containing 10 to 20-fold more L-2HG than most tissues^10,19^. However, the specific cell stage in which 2HG is accumulated and whether or not it regulates testicular cell biology is an enigma. Thus, we sought to delineate the levels of 2HG in different stages of germ cell development and assess whether and how it is involved in germ cell biology. We found that 2HG is generated only at specific stages of the spermatocyte lineage and its function is not related to the inhibition of DNA/histone demethylation, but rather to its effects on the morphology of centromeres and heterochromatin confined to chromocenters at these specific stages. These effects are necessary for the proper progression of meiosis and thus directly link 2HG to controlling cell division.

## Results

### Stra8-Tom mice – a tool for isolating viable germ cell populations from adult testis

To fluorescently label the male germ cell lineage we crossed Stra8-cre mice, that express Cre recombinase in early stage spermatogonia^20^ with CAG-LSL-tdTomato mice to generate “Stra8-Tom” mice. As expected, tomato expression was only observed within lumens of the seminiferous tubules and there was no overlapping of tdTomato expression with the Sertoli cell marker SOX9, confirming the expected germ cell lineage restriction (**Figure S1A**). Unexpectedly, tomato expression significantly decreased along the germ lineage differentiation stages (**Figure S1B**). This phenomenon likely occurs due to decreasing activity of the CAG promoter along the different stages, as ROSA driven fluorescence (via crossing of ROSA-LSL-YFP with Stra8-cre mice) was uniformly distributed (**Figure S1C**). We noted that this gradual decrease could provide a means for isolating large numbers of germ cells at different stages of the germ cell lineage. FACS analysis of cell suspensions prepared from the testes of Stra8-Tom mice after dead cell removal (Forward scatter A vs. red fluorescence) revealed 6 discrete cell populations (**Figure 1A**) owing to the stepwise decrease of tdTomato levels and cell size differences.

**Figure 1:**
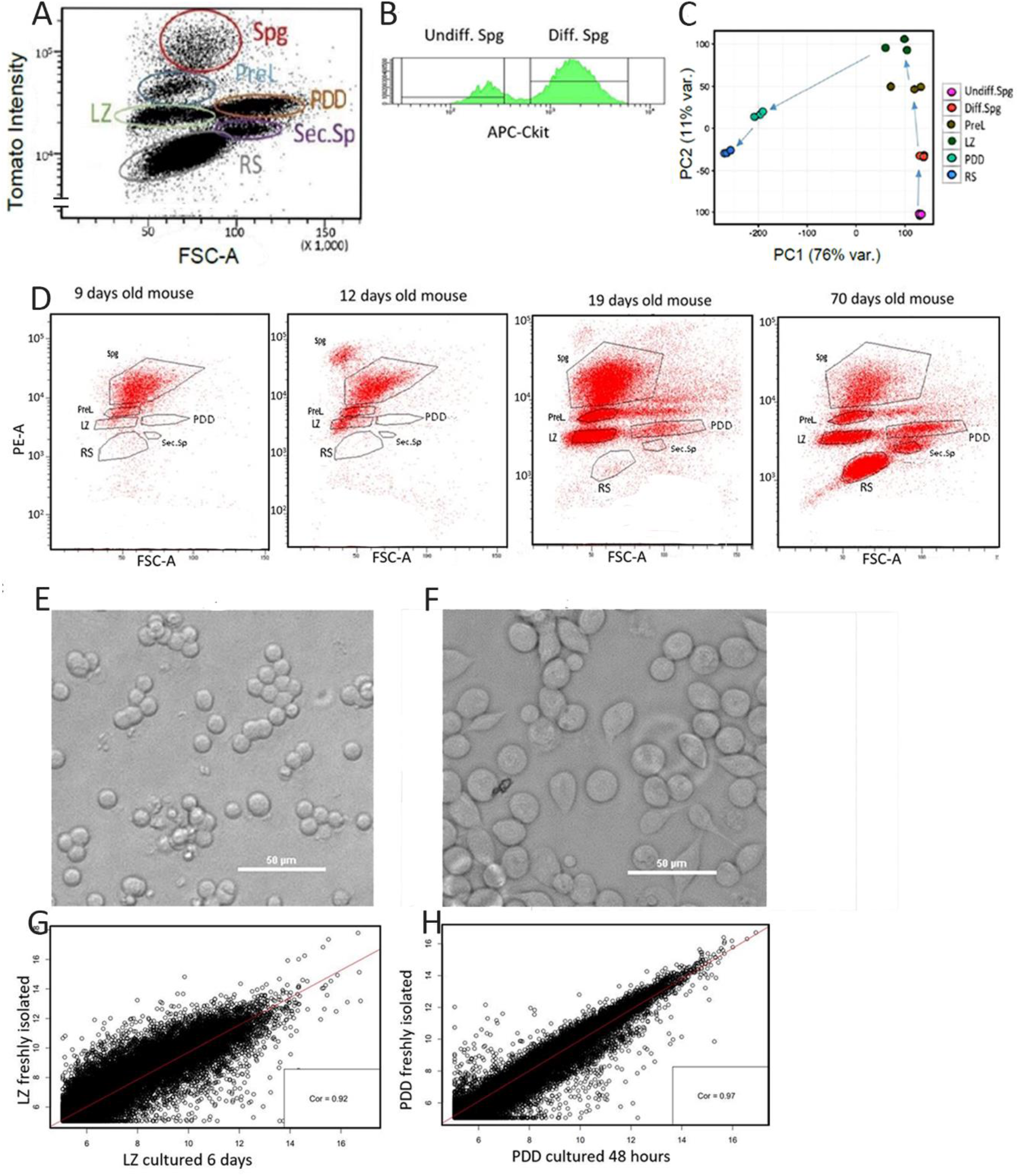
Isolating highly purified populations of germ cells with Stra8-Tom mice: **A.** Representative FACS analysis of Stra8-Tom testis cell suspension. X-forward scatter; Y-red fluorescence. **B**. FACS analysis of Spg cells stained with APC conjugated Ckit antibody. **C.** PCA plot of mRNAseq data from the indicated populations (n=3 for each population). **D**. FACS analyses of testicular cell suspensions from the indicated ages-more differentiated cell populations gradually appeared with age. **E-F** LZ and PD cells were grown in culture for 6 days and 48 hours, respectively. Shown are representative phase contrast images. The notable change over time was that LZ cells tend to form clusters. Scale bar: 50 μM. **G**. Gene expression (RNAseq) from freshly isolated LZ cells vs. LZ cells cultured for 6 days. Pearson’s product-moment correlation: r=0.92; p<2X10^−16^. **H**. Gene expression (RNAseq) from freshly isolated PDD cells vs. PDD cells cultured for 48 hours. Pearson’s product-moment correlation: r=0.97; p<2X10^−1^

To assess whether each population represents specific stages of the germ cell lineage and their cell purity, we assessed DNA content with Hoechst or propidium iodide (**Figure S1D**) coupled with immunostaining for known markers of different stages of the male germ cell lineage including DMRT1, PLZF, SCP3 and γH2AX (**Figures S1E-S1H**). Immunostaining for cKit further separated between undifferentiated and differentiating spermatogonia (**Figure 1B**). Our analysis revealed that each of the 7 cell populations (including spermatogonia separated with cKit antibody) was highly enriched for a specific stage of the germ cell lineage. Thus, we were able to sort undifferentiated (Undiff) and differentiating (Diff) spermatogonia (Spg), preleptotene (PreL), leptotene/zygotene (LZ), pachytene/diplotene/diakinesis (PDD), secondary spermatocytes (Sec.Sp) and round spermatid (RS) populations. The composition and purity of each population is detailed in **Figure S1**. Next, we performed mRNA sequencing analysis on triplicate samples from the different cell populations (GSE162740). The secondary spermatocyte population was not analyzed due to cell number limitations. Unsupervised principle component analysis (PCA) revealed that each stage clusters separately and has ordered the cells along the known differentiation program (**Figure 1C**). The largest change in gene expression (PC1) was observed between LZ and PDD stages, both belonging to the meiosis I prophase. Nearly 10,000 genes showed more than 2-fold change in gene expression between these two stages (**Figure S1I).**

To further validate our method for separating germ cell populations, we harvested testicular cells from Stra8-Tom mice of the indicated ages and subjected them to FACS analysis. Newborn male mouse testes harbor only spermatogonial stem cells; later stages gradually appear in a semi-synchronous temporal pattern until mice reach the first round of sperm differentiation. Thus, more differentiated cell populations gradually appeared with age, as expected (**Figure 1D**). Next, we optimized culture conditions that allowed us to culture LZ cells for up to 10 days (**Figure 1E**) and PDD cells for up to 3 days (**Figure 1F**) with minimal cell death. We subjected mRNA isolated from freshly sorted or cultured LZ and PDD cells to mRNAseq analysis (GSE169014). Transcriptome comparison revealed that the cell populations maintain their transcriptional landscape throughout the culture period (r = 0.92 and 0.97 for LZ and PDD populations, respectively, **Figure 1G-H**). Thus, in addition to confirming the utility of the cultured cells to probe germ-lineage cell biology this suggests that the transcriptional landscape of these stages is stable in these *in vitro* conditions and that progression along the differentiation program may require extracellular signals.

### L-2HG is produced in pachytene/diplotene/diakinesis cells by LDHC

Previous studies showed that whole testis extract harbors a high content of 2HG, yet the specific cell stages that generate this metabolite were not defined^10,19^. As Stra8-Tom mice provide us an opportunity to isolate sufficient numbers of viable cells for biochemical analyses, we sought to define the abundance of 2HG in different stages of the germ cell lineage. Thus, we analyzed extracts of metabolites from 4 major cell populations using LC-MS. This revealed that the content of 2HG increases 12-17-fold in PDD and RS populations as compared to LZ and Spg cells (**Figure 2A**). Chiral derivatization combined with LC-MS confirmed that 2HG levels are higher in PDD and RS cells vs. earlier stages, and showed that L-2HG is the predominant enantiomer while D-2HG content is negligible (**Figure S2A**).

**Figure 2:**
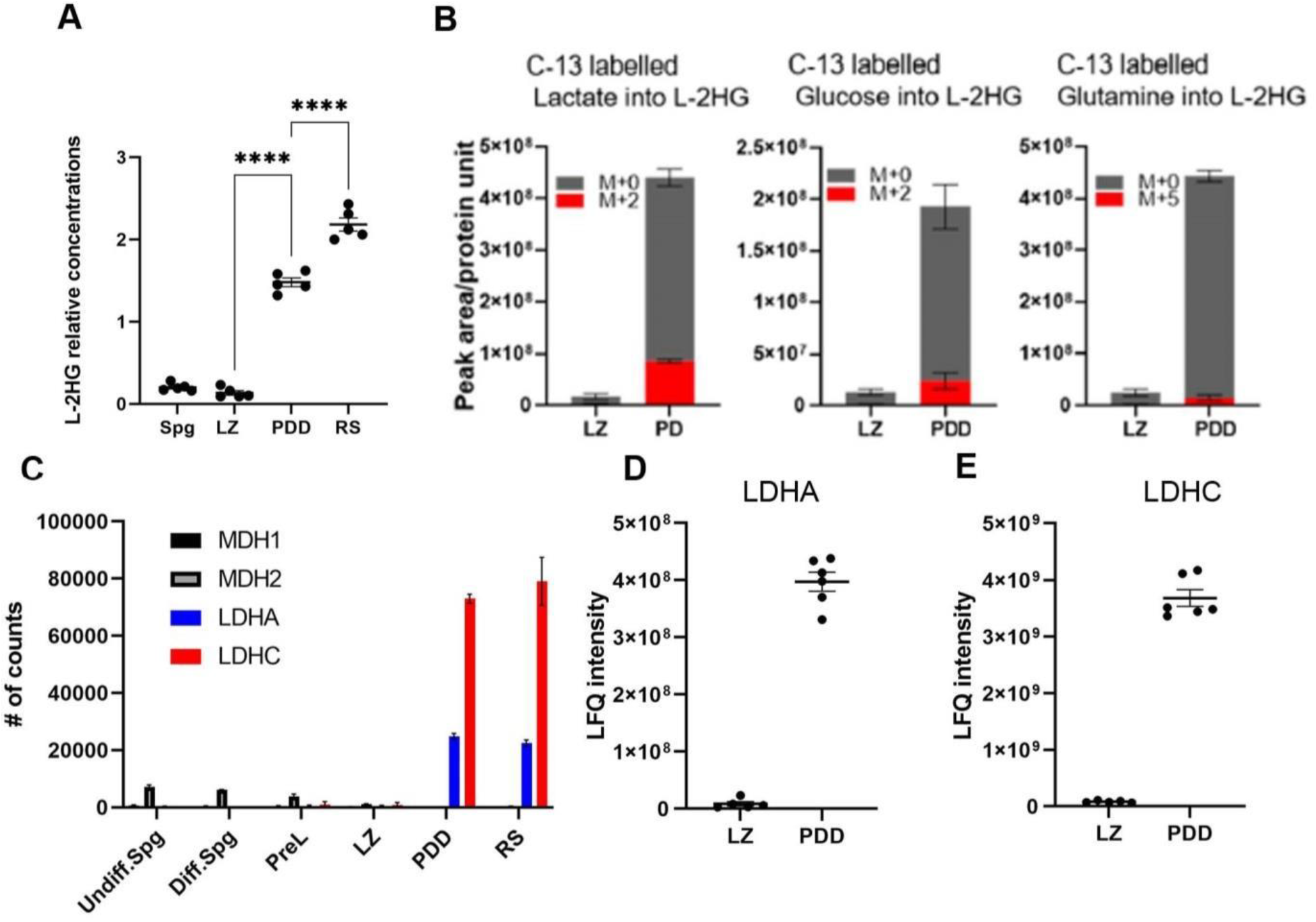
2HG is generated by LDHC in PDD and RS cells: **A.** 2HG content (in the indicated populations) was measured by LC-MS in 5 independent experiments. Results are corrected for 10^6^ cells and for cell volume. Cell volume measurements are provided in **Table S1**. Mean±SE; One-way Anova-Tukey’s multiple comparisons. **B.** Incorporation of 13C-lactate-3, 13C-glucose-6 and 13C-glutamine-5 into 2HG isotopologues was measured using LC-MS. Peak area corrected to protein content after 2 hrs of incubation of LZ and PDD cells. Mean±SE of 4, 5 and 3 independent experiments of the tracing of lactate, glucose and glutamine, respectively. **C.** Expression values (counts) of LDHA, LDHC, MDH1 and MDH2 derived from mRNAseq data, mean±SE of 3 samples. **D-E**. Relative content of LDHA (**D**) and LDHC (**E**) derived from MS data. 5 and 6 independent samples of LZ and PDD cells, respectively, mean±SE.

To assess the carbon source for L-2HG we incubated LZ and PDD cells for 2 hours with 13C lactate, 13C glucose and 13C glutamine and measured incorporation of the tracer into 2HG. As expected the pool of L-2HG was about 25-fold higher in PDD compared to LZ cells. 2 hours after incubating cells with labeled lactate, 20% of the L-2HG pool in PDD cells was labeled and appeared as the M+2 isotopologue, whereas only negligible amounts of labeling was detected in LZ cells. Incubation with 13C glucose labelled about 14% of the pool, while there was almost no incorporation of 13C carbons from 13C glutamine. This suggests that L-2HG is constantly generated in PDD cells while LZ cells do not actively generate L-2HG and that lactate is a main source for L-2HG generation (**Figure 2B**).

L-2HG can be produced by several enzymes including the three isoforms of lactate dehydrogenase (LDHA, LDHB, LDHC) and the two isoforms of malate dehydrogenase (MDH1, MDH2). Inspection of the mRNAseq data revealed that LDHA and the testis specific LDHC, are markedly upregulated in PDD compared to LZ cells, while LDHB is not expressed in either population. MDH1 and MDH2 are expressed at low levels and show decreased expression along differentiation (**Figure 2C**). In line with the mRNA expression data, mass spectrometry of protein extracts revealed that both LDHA and LDHC are markedly overexpressed in PDD vs. LZ cells (**Figure 2D-E**), thus theoretically both isoenzymes of LDH could synthesize L-2HG. Previously, it was shown that whole testis extracts from LDHC null mice^10^ do not synthesize L-2HG and contain very low levels of L-2HG. Taken together, we concluded that LDHC, that is expressed in the germ lineage beginning at the pachytene stage is the enzyme responsible for L-2HG production in PDD and RS populations.

### LDHC localizes along chromosomes and in centromeres of PDD cells

Several studies reported that certain cytoplasmic and mitochondrial metabolic enzymes are transported to the nucleus where they can generate metabolites that may be involved in genome and epigenome regulation^21–26^. As 2HG is thought to act as a chromatin modifier we studied the localization of LDHC by performing immunofluorescent staining. As expected, there was a clear staining of LDHC in PDD but not in LZ cells. Remarkably LDHC was distributed in several cell compartments. While significant staining was observed in the cytoplasm of PDD cells, it was also detected in the nuclei, where its distribution changed between the PD and diakinesis stages. We discovered that in PD cells, LDHC was localized along the chromosome axes as evident by colocalization with SCP3 (**Fig 3A-B**) and was particularly intense in the centromere region.

**Figure 3:**
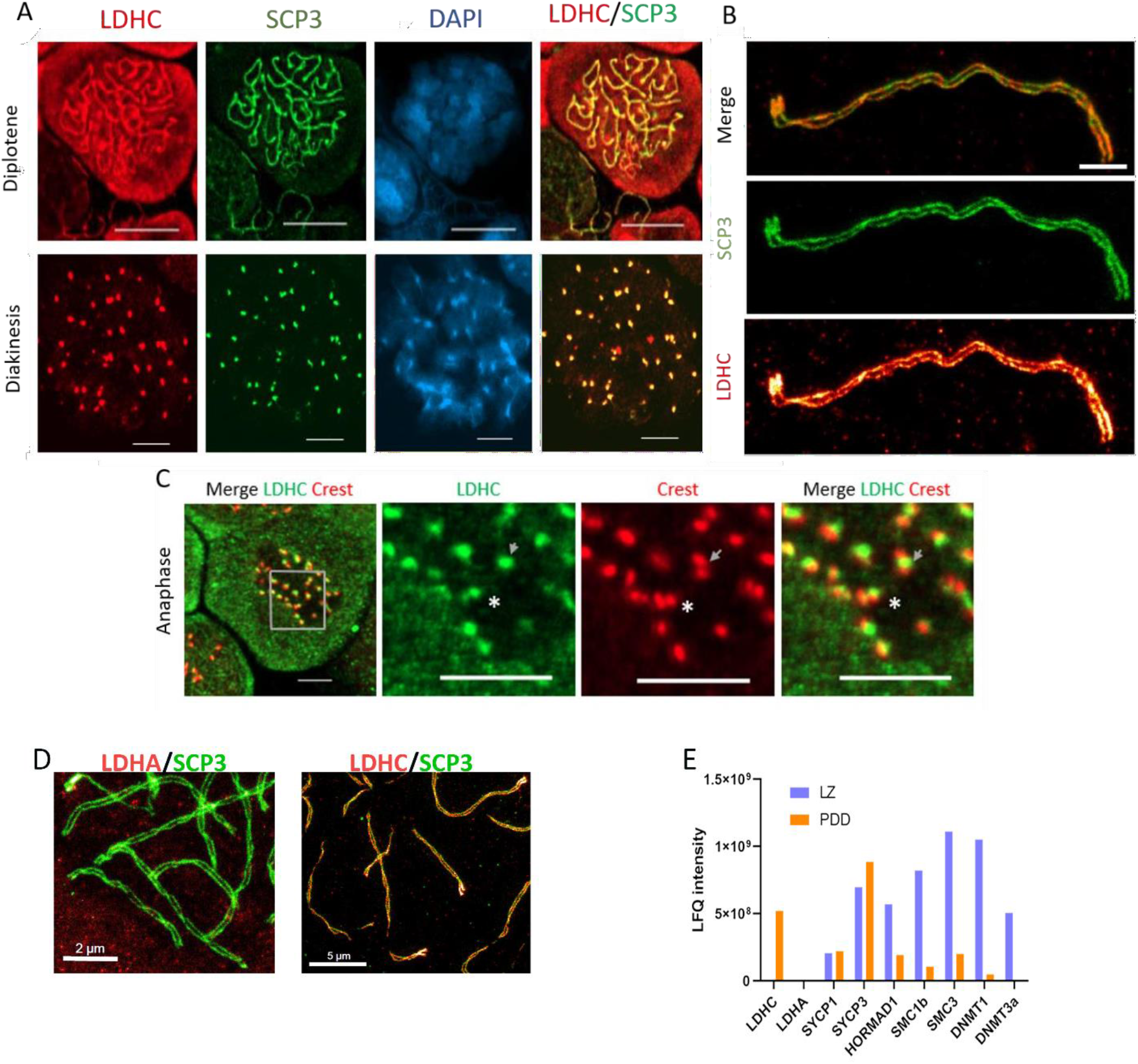
LDHC localizes along chromosomes and in centromeres: **A-C** Immunostaining of freshly isolated testicular cells for LDHC (199891-A antibody). Cells were identified using SCP3 (A and B) or CREST (C) antibodies. **A.** Confocal microscope image of diplotene and diakinesis cells, scale bar 10 μm. **B.** STED image of pachytene chromosome (released from freshly isolated cells), scale bar 1μm. **C.** Confocal microscope image of anaphase cell, scale bars: 5 μm (left, whole cell) and 1μm (right, enlarged insert). **D.** STED image of pachytene chromosome stained for LDHA (left, scale bar 2μm) and LDHC (right, scale bar 5μm) and SCP3, **E.** LFQ intensities of chromatin bound proteins obtained from MS data. Proteins bound to chromatin were extracted with high salt solution. Note high content of LDHC in PDD population as compared to some proteins established as bound to meiotic/mitotic chromosomes in literature^27,28^. LDHA was non-detectable. One of two experiments is shown.

This pattern was confirmed using high resolution stimulated emission depletion (STED) microscopy (**Figure 3B**). This staining pattern was also observed using a different anti-LDHC antibody which targets a different LDHC-epitope and when staining nuclear spreads^29^ (**Figure S2B-C**. LDHC staining of centromeres was also prominent in the diakinesis stage (**Figure 3A**) and was adjacent to centromeres in anaphase (**Figure 3C**). Of note, LDHA staining was also noted in the nuclei of PD cells, but unlike LDHC was not localized to chromosomes (**Figure 3D** and **Figure S2D**). To verify the presence of LDHC on chromatin we extracted chromatin-bound proteins from LZ and PDD cells and submitted them for mass spectrometry analysis. This analysis confirmed the presence of LDHC in the chromatin bound fraction in PDD but not LZ cells. Notably LDHA was not detectable in extracts from both cell types (**Figure 3E**). Thus, our findings reveal that L-2HG is generated in PDD stages by LDHC and suggest that L-2HG may be generated in PDD cells nuclei along the chromosomes, particularly in centromeric regions and solely in the centromeres of the diakinesis stage. The putative high local concentrations of L-2HG in centromeric regions in PDD cells led us to hypothesize that it may control centromere function.

### L-2HG and LDHC guard the morphology of centromeres in diakinesis

Centromeres are highly dynamic structures, uniquely changing their morphology to allow their proper function in cell division^30,31^. In order to study the function of L-2HG we sought to modulate its levels using oxamate, a known pan-LDH inhibitor^32^. Incubation of PDD cells with oxamate for 48 hours decreased the content of L-2HG by 10-fold (**Figure 4A**). Next, we assessed the effect of oxamate treatment on centromeres in diakinesis cells, in which centromeres are fully mature to facilitate the first meiotic division. Oxamate treatment, both *in vitro* and *in vivo*, caused significant enlargement of centromere area (measured using CREST antibodies) in diakinesis cells (**Figure 4B-C**). As oxamate may have other effects, in addition to L-2HG depletion, we performed a ‘rescue’ experiment with exogenous L-2HG. The addition of cell-permeable octyl L-2HG together with oxamate for 12 hours markedly diminished the oxamate induced centromere de condensation (**Figure 4C**). This effect was observed already at 6 hours (data not shown). To assess the ability of 2HG to affect centromere condensation in even shorter time frames, we tested whether adding 2HG to cells isolated from oxamate treated mice (in which centromeres are already enlarged) can restore normal centromere size.

**Figure 4:**
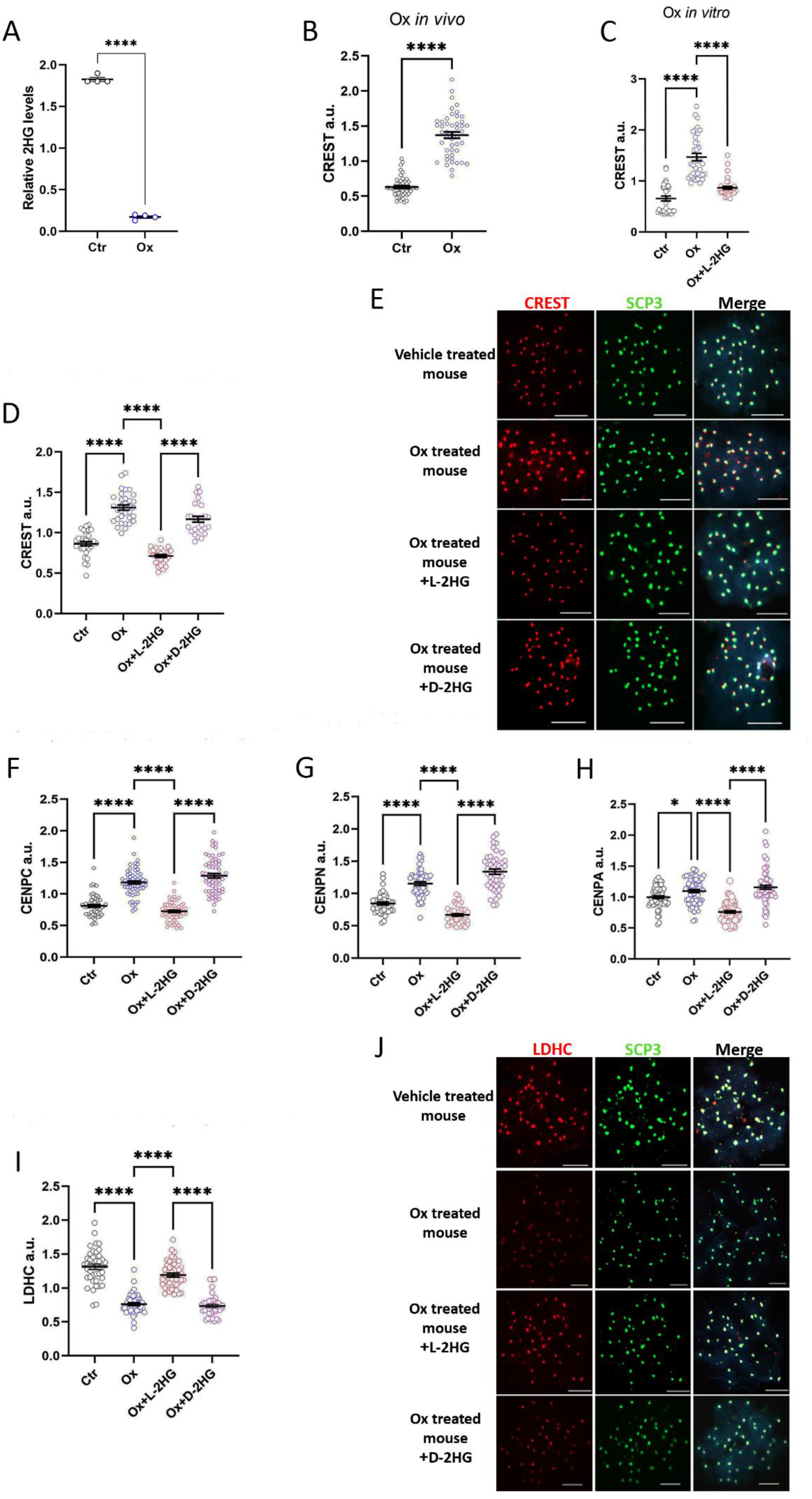
L-2HG and LDHC guard centromere morphology in diakinesis cells. **A**. Isolated PDD cells were incubated for 48 hrs with vehicle or 24 mM oxamate, relative concentrations of 2HG were measured using LC-MS. Mean ±SEof 4 experiments, t-test. **B-C**. Cross sectional area of centromeres of diakinesis cells measured with CREST antibodies staining. Diakinesis cells were isolated either from from mice injected once with oxamate (1.3g/kg, sacrificed after 24 hrs) -**B**, or from diakinesis cells incubated with 24 mM oxamate or with 24 mM oxamate and 0.3 mM permeable L-2HG for 12 hrs in culture-**C**. Nuclear spreads were prepared and stained with CREST. Each data point represents the mean area of all centromeres of one cell. Mean±SE of 30-50 cells in each treatment, **B** Mean ±SE t-Test, **C** Mean ±SE Tukey’s multiple comparisons test. **D-J**. Diakinesis cells were isolated from vehicle or oxamate treated mice (as in **B**). Cells from oxamate injected mice were incubated with 0.6 mM cell permeable 2HG (either L or D) for 10 min at 32°C. Nuclear spreads were stained with the following antibodies: CREST (**D, E**), CENPC (**F**), CENPN (**G),** CENPA (**H**) or LDHC (**I-J**). Each data point represents mean area of all centromeres of one cell. Mean±SE of 30-60 cells in each treatment; Tukey’s multiple comparisons test. Stained areas were measured using Zeiss ZEN 3.3 software and expressed as fraction of the mean of all experimental groups in arbitrary units (a.u). **E**. Representative pictures of centromere area stained with CREST and quantified in **D**. **J**. representative pictures of centromere area stained with LDHC and quantified in **I**. Scale bars in **E,J**: 10 μm.

Remarkably, even 10 min incubation of cells with octyl-L-2HG at 32°C, just before chromosome-spread preparation and fixation with paraformaldehyde, sufficed to revert centromere size. Similar incubation with the octyl-D-2HG enantiomer had no such effect (**Figure 4D, E**), indicating a stereospecific effect. CREST antibodies recognize mainly CENP-A, CENP-B and CENP-C, components of core centromeric nucleosome complex (CCNC)^33^. In order to define components that undergo de compaction we analyzed staining with antibodies against CENP-C, CENP-N and CENP-A. CENP-C contains intrinsically disordered regions (IDRs)^34^, thus it can be prone to liquid-liquid phase separation. CENP-N was recently shown to promote centromeric chromatin compaction in human cell lines ^35^. In this context it was interesting to observe that both CENP-C and CENP-N staining showed identical pattern of changes to those shown with CREST antibodies (**Figure 4 F,G**). This pattern was not observed with CENP-A immunostaining (**Figure 4H**). Oxamate treatment enlarged CENP-A area only slightly by 1.1 fold as compared to 1.45 and 1.37 fold for CENP-C and CENP-N respectively. This can be expected since CENP-A forms centromere-specific nucleosome at the base of the CCNC to which CENP-C, CENP-N and other proteins bind. The effect of oxamate was reversed by exogenously added L but not D-2HG.

Remarkably, oxamate treatment resulted in LDHC detachments from the centromeres, an effect which was rapidly restored by adding L-2HG (**Figure 4 I,J).** Thus, de compaction is associated with deposition of LDHC from centromeres and the repositioning of this enzyme depends on L-2HG.

### L-2HG guards spatial organization of centromeres and chromocenters of diplotene cells

To assess the effects of 2HG on centromeres of diplotene cells, we incubated cells for 24-48 hours with oxamate (to decrease L-2HG content) or with oxamate together with cell permeable 2HG.Immunostaining with CREST antibodies, revealed that oxamate treatment caused expansion of the centromeres’ cross-sectional area by 1.8-fold after 48 hours of treatment (**Figure 5A and Figure S**3**A**). Oxamate induced centromere decondensation was completely prevented by addingoctyl-L2HG to the incubation medium, suggesting that L-2HG controls centromere condensation in diplotene cells, similar to its role in diakinesis cells. To obtain high resolution images of this change we used STED microscopy. High resolution imaging confirmed the effect on centromere cross sectional area and revealed that depleting L-2HG levels resulted in a diffuse pattern of CREST staining, contrasting the condensed pattern observed in the presence of physiological levels of L-2HG in diplotene cells (**Figure 5B**). The pericentromeric region, also known as the chromocenter, is heterochromatinized and silenced by the repressive histone mark H3K9me3 which is bound by the silencing protein HP1α. Chromocenters are particularly prominent in the diplotene stage before diakinesis, allowing us to study the effects of 2HG on chromocenters and the centromeres embedded within them, as well as to assess possible interactions between these two nuclear subcompartments.

**Figure 5:**
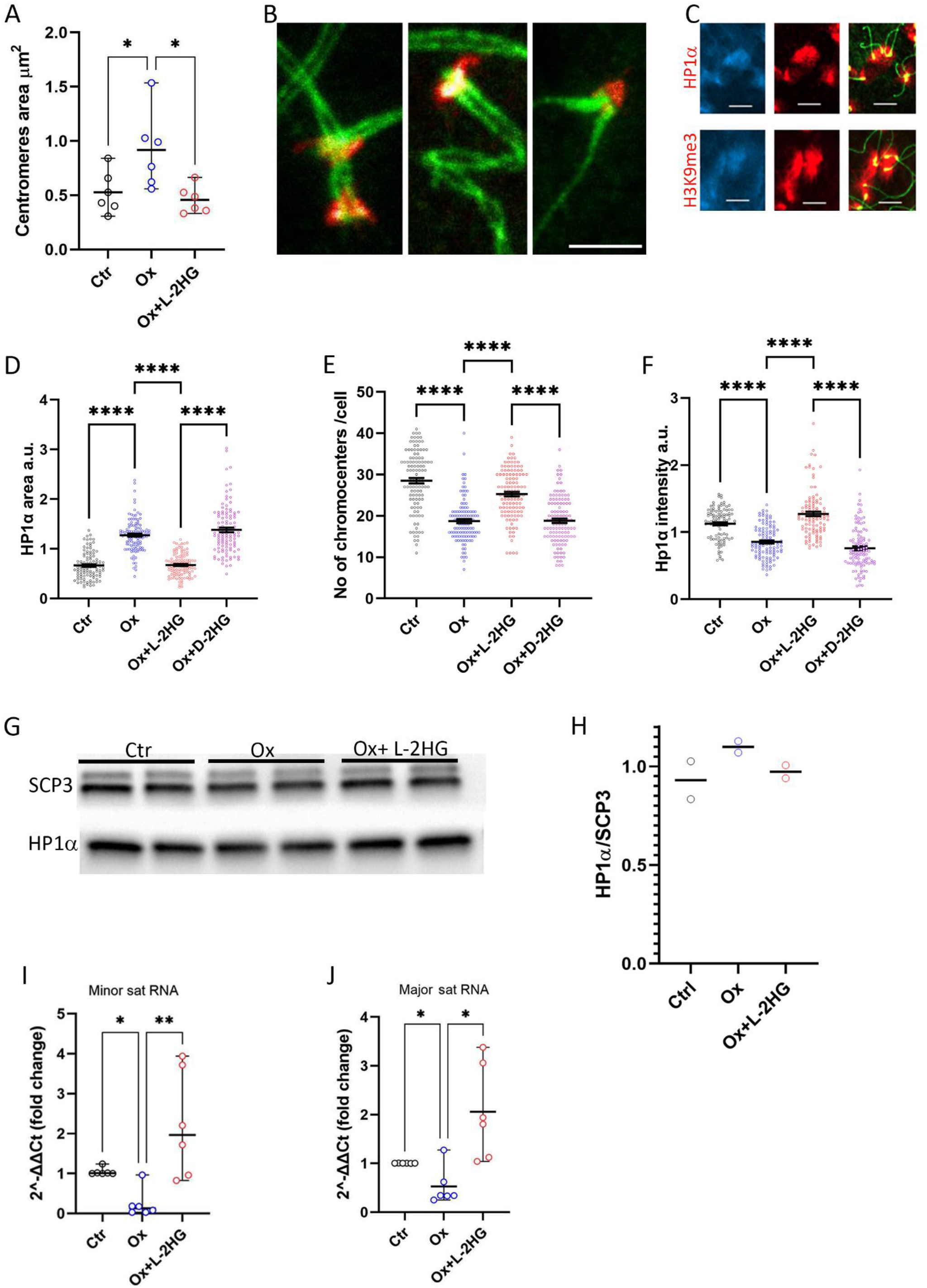
L-2HG guards spatial organization of chromocenters in diplotene cells. **A-B, D-L**. Isolated PDD cells were incubated for 24 (D-H,) or 48 hrs (A-B, I-J) with vehicle, 24 mM oxamate or 24 mM oxamate with 0.3 mM cell-permeable L/D-2HG, as indicated. **A**. Centromere area was measured on CREST-stained nuclear spreads using NIS elements analyser software. Each data point represents the mean of 200 to 400 regions of interest measured in 6 different experiments, median with range, Bonferroni correction for multiple comparisons, one tail **B.** Representative high-resolution STED image, of nuclear spreads stained with SCP3 (green) and CREST (red), scale bar 1 μm. **C.** HP1α and H3K9me3 (red) demarcate chromocenters (DAPI-blue). Diplotene chromosomes stained with SCP3 (green). Scale bar 5 μm. **D**. Areas of chromocenters of diplotene cells were calculated using NIS elements software and expressed as a fraction of the mean of all experimental groups in arbitrary units (a.u). Each data point represents the total area of one cell stained with HP1α antibody, mean±SE of about 100 cells in each group, one-way Anova, Tukey’s multiple comparisons. **E**. Numbers of chromocenters were counted for each diplotene cell using data from D. **F**. Intensity of red fluorescent stain of HP1α/area unit/cell. Mean±SE of about 100 cells in each group. **G-H**. Immunoblot analysis for HP1α content in whole cell protein extracts of PDD cells normalized to SCP3, one of two representative experiment. **I-J**. qPCR performed for minor (**I**) or major (**J**) satellite RNA; median with range of 6 experiments, Bonferroni correction for multiple comparisons, one tail.

Immunostaining for either HP1α or H3K9me3 clearly demarcates DAPI-intense chromocenters (**Figure 5C**). Diplotene cells incubated for 24 hrs with oxamate showed enlargement of chromocenter area which was observed with both HP1α (**Figure 5D**) and H3K9me3 (**Figure S3B**) staining, 1.9 and 1’.6 fold, respectively. This phenotype was rescued by L-2HG and was enantiomer specific. Since chromocenter clustering is observed in germ cells at the meiotic prophase and plays an important role in subsequent meiotic processes^36^ we counted the number of chromocenters in each treatment group (**Figure 5E**). We found that the number of chromocenters decreased with oxamate treatment by 1.5-fold implying that this treatment increased chromocenter clustering. This phenotype is rescued by L-, but not D-2HG indicating that L-2HG regulates chromocenter area and clustering in diplotene cells. The enlargement of heterochromatin area upon L-2HG depletion could be mediated either by an expansion of heterochromatization to lateral nucleosomes or by decreased compaction of pre-existing HP1α/H3K9me3 marked nucleosomes. To resolve these two options, we measured Hp1α staining intensity in chromocenters. We detected a 1.3-fold decrease, for HP1α staining intensity under oxamate treatment (**Figure 5F**). In addition, we did not observe a significant change in total HP1α levels in the different treatment groups (**Figure 5G-H**).

Measuring the global levels of multiple histone modifications using a comprehensive ELISA KIT did not reveal reproducible changes in the global levels of any modification (**Figure S3C**). Thus, L-2HG inhibits chromocenter clustering and de compaction. The pericentromeric and centromeric regions are typically gene-poor, with little associated mRNA transcription due to the presence of constitutive heterochromatin marks. However, the satellite sequences in these regions are transcribed into satellite RNAs. Satellite RNA expression is known to be regulated along the cell cycle and satellite RNAs were shown to be important for centromere function^37,38^. We hypothesized that L-2HG could affect satellite RNA expression. Indeed, expression of both minor and major transcripts significantly decreased, 10 and 3-fold respectively, in oxamate treated PDD cells. Adding L-2HG completely rescued this phenotype (**Figures 5I-J**). In accordance, it was recently shown that UV induced heterochromatin de compaction in mitotic mouse embryonic fibroblasts similarly reduced satellite RNA expression^39^.

### 2HG is necessary for normal progression of meiosis

Centromeres normally separate from each other in the diakinesis stage. It was previously reported that on average four centromeres remain paired in the diakinesis stage of male meiosis^40,41^. When examining diakinesis cells isolated from untreated mice we noted an average of two unseparated pairs of centromeres in line with previous reports; centromere pair numbers increased approximately 2-fold after oxamate treatment (**Figure 6A-B**). This effect could also be observed in cultured diakinesis cells treated with oxamate for 24 hours and was prevented by the addition of L-2HG together with oxamate (**Figure S3D-E**).

**Figure 6:**
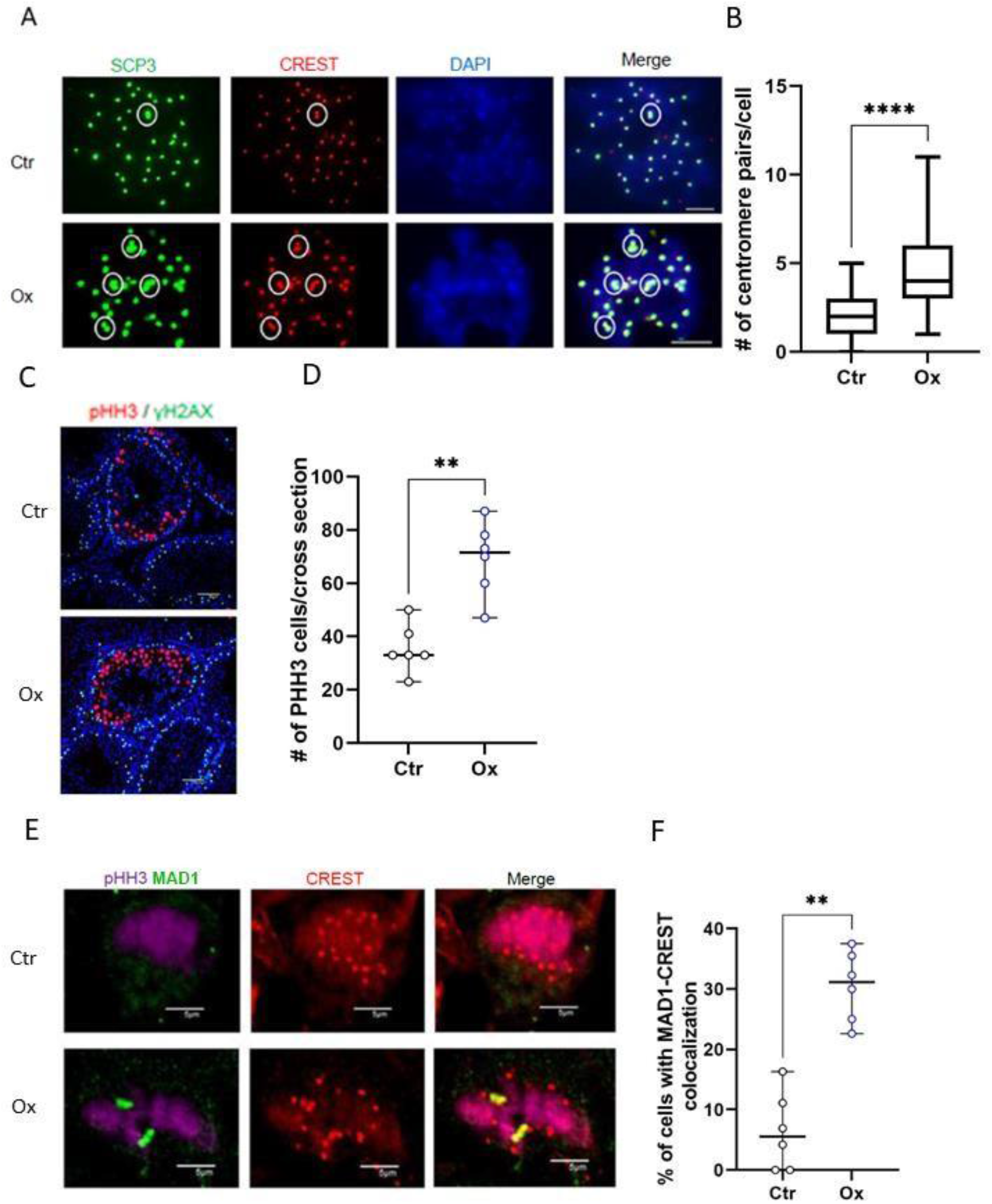
Oxamate treatment causes meiotic centromere dysfunction and an activation of the spindle assembly checkpoint *in vivo*. **A-B**. Mice were injected once with oxamate 1.3g/Kg or vehicle. Diakinesis cells present in the isolated PDD population were stained with SCP3 and Crest. **A.** Centromere pairs are marked with white circles, scale bar 10 μm. **B.** Number of centromere pairs were counted in 40 cells in diakinesis, median with range, Mann-Whitney. **C-F**. Mice were treated with daily oxamate injections (as above), or vehicle for 7 days (n=6). Paraffin sections were stained with pHH3, γH2AX and counterstained with DAPI. Representative image, scale bar 10 μm (**C**) and quantification of pHH3 positive cells (NIS elements analyser software) (**D**). One experiment out of two is shown. Median of 6 mice with range. Mann Whitney. **E**-**F** Paraffin sections were stained with PHH3 (purple), MAD1 (green) and Crest (red). pHH3 positive cells were assessed for the presence of overlapping staining for MAD1 and CREST. Representative image, scale bar 5 μm (**E**) and the percent of MAD1-CREST overlapping cells from pHH3 positive cells; at least 30 nuclei per mouse. Each data point represents a single mouse. 6 mice in each group; median with range; Mann Whitney (**F**).

The above findings prompted us to study whether 2HG regulates centromere functions in PDD cells *in vivo*. First, we checked whether oxamate treatment *in vivo* recapitulates effects noted in the *in vitro* setting in PDD cells. To this end we treated mice with oxamate for 24 or 48 hours and analyzed its effect on centromere diameter, chromocenters (H3K9me3 staining), and satellite RNA expression. Notably, it is not possible to perform rescue experiments with cell permeable L-2HG in the *in vivo* setting. Oxamate treatment increased average centromere cross sectional area of diplotene cells by 2.6-fold in treated mice (**Figure S4A**). Chromocenters area demarcated by H3K9me3 increased 1.6 fold, while the average number of chromocenters and staining intensity/area unit decreased, 1.4 and 3.5 fold, respectively (**Figures S4B-D**). Oxamate also reduced expression levels of minor and major satellite RNAs in PDD cells by about 5 and 2.5-fold, (**Figures S4E** and **S4F,** respectively). Thus, all the effects of oxamate treatment on centromere and chromocenter morphology *in vitro* were recapitulated in the *in vivo* setting, allowing us to test whether L-2HG is necessary for the proper function of centromeres within a living mouse.

The meiotic prophase in male mice is extremely long, lasting about 2 weeks, compared with a 30-60 minutes duration of the mitotic prophase; the pachytene-diplotene stages of mouse male meiosis last about one week^42,43^. Therefore, to assess the effect of L-2HG depletion on meiosis progression *in vivo* we treated mice with oxamate for 7 days. Control mice were treated with vehicle and mice were euthanized on day 8 of the experiment. Histological analysis of PAS stained slides and immunostaining for cleaved caspase 3 did not reveal signs of toxicity (**Figure S5**). Next, we stained testis sections with antibodies against histone H3 phospho-serine 10 (pHH3) - a marker of cells in late G2 and M phase. Oxamate treatment resulted in a 2-fold increase in the number of pHH3 positive cells (**Figures 6C-D**). This could either suggest an increased proliferation rate or cell cycle arrest, as H3 remains phosphorylated in cells that are arrested in the G2M checkpoint. Cell cycle arrest due to centromere malfunction is executed by activation of the spindle assembly checkpoint (SAC).

Thus, we co-stained testis sections with pHH3, MAD1 and CREST antibodies and quantified the number of pHH3 positive cells harboring MAD1 localized to the centromeres – an indication for SAC activation^44,45^. Remarkably, oxamate treatment increased the number of cells in which the SAC is activated 6-fold (**Figures 6E-F**), supporting a role for L-2HG in ensuring the fidelity of centromere function in male meiosis. Taken together, our data suggests that the high physiological levels of L-2HG present in PDD cells are necessary for proper assembly of the centromeric complex and its absence could result in centromere de-condensation and malfunction during meiosis in the male germline.

## Discussion

The oncometabolite 2HG gathered attention when it was discovered that it is highly abundant in tumors harboring activating mutations in IDH enzymes. It is also upregulated in lymphocytes in specific states and in hypoxic cells where it governs cellular transcriptional responses^15,16,18^. While L-2HG was known to be most abundant in the testis^10,19,46^, its distribution along the germ cell lineage and whether it accumulates as a metabolic byproduct or plays an active role in regulating germ cell biology remained enigmatic. We developed a flow cytometry-based method for isolating and culturing high numbers of viable germ cells of specific stages, similar to a previously published method reported by Page and colleagues^47^; however, our method does not require retinoic acid-based synchronization. Using this method, we revealed that the expression of 2HG producing enzymes is restricted to specific stages of the first meiotic prophase and only initiates after completion of the zygotene stage. Accordingly, we detected high levels of L-2HG in PDD and RS stages, but not in earlier stages. We have confirmed that LDH, is the major source of L-2HG in the testis, by using the pan-LDH inhibitor oxamate. In accordance,it was previously shown that LDHC null mice have low levels of L-2HG in the testis^10^. Remarkably, we now revealed that LDHC is markedly upregulated at the PDD stage and is associated with centromeres of PDD cells and along chromosomes in the PD stages. While we cannot measure local concentrations of L-2HG in the centromere region we speculate that the unique distribution of LDHC could generate high concentrations of L-2HG in these subnuclear compartments. As LDHC null mice have low levels of 2HG in the testis, it seems that LDHA is not sufficient to produce 2HG in this tissue. Still, LDHA is also highly expressed in PDD cells, but not in earlier stages. While LDHA is also detected in the nuclei, it does not localize to centromeres or chromosomes.

We found that the expression of minor and major satellite RNAs expressed from pericentromeric regions depends on L-2HG. Satellite RNAs allow for proper centromere function during mitosis and meiosis, possibly through maintaining the compaction of the centromere either via tethering or recruiting centromeric proteins, or through induction of phase separation processes, similar to other long noncoding RNAs^30,37,38,48^. It was shown that both overexpression and under-expression of satellite RNAs cause defects in centromere morphology and chromosomes segregation in mitotic cells^49,50^. Recently, satellite RNAs were also shown to control the fidelity of meiosis^38,51^. The way by which L-2HG controls the expression levels of satellite RNAs is not yet clear. This could be via classic transcriptional regulation of the satellite RNA genes, possibly by controlling local DNA or histone methylation. It is also plausible that the changes in the total levels of satellite RNAs reflect downstream effects of heterochromatin condensation (as indicated by H3K9me3 and HP1α staining), which could alter their transcription or degradation rates. As pointed above, our results are very similar to those described for de compacted chromocenters of embryonal mouse fibroblasts where transcriptional silencing was also prominent^39^.

It is widely believed that the regulatory roles of L- and D-2HG in cancer and physiology are mediated through inhibition of αKG dependent demethylases^15,16^. However, the enantiomer specific rapid (10 min incubation) condensation of diakinesis centromeres by L-2HG, occurring at 32°C, supports the possibility that L-2HG induces a conformational change in a specific target protein, via binding of an allosteric regulatory site according to a lock and key mechanism. Notably, the few studies that compared the inhibitory activity of L- and D-2HG towards different histone demethylases did not observe marked differences between the two enantiomers^15,16^, in line with our hypothesis that the physiological roles of L-2HG in the testis are not mediated solely by inhibiting DNA and histone demethylases. Interestingly, phosphorylation of a structured region of HP1α, a key effector of heterochromatinization, was shown to expose a disordered region in this protein to form homotypic polymer-polymer phase separated (PPPS) chains and affect heterochromatin structure^52^. We hypothesize that a similar effect occurs upon binding of L-2HG to a specific target protein. Our data show LDHC localization to the centromere requires normal levels of L-2HG raising the possibility that this protein could be the L-2HG binding protein. We hypothesize that the conformational change induced by binding to L-2HG renders the target protein more capable of forming high order structures resulting in condensation of the chromocenter and centromere (**Figure 7**). To our knowledge, the regulation of centromeres compaction by a metabolite that is synthetized at the specific stage of germ cells differentiation is rather unique, however it is possible that similar metabolic effectors will be found in other systems. Identifying target protein/s, whose conformation is regulated by L-2HG could reveal if the mechanism we discovered in the male germline may be operative in other physiological or pathological conditions.

**Figure 7:**
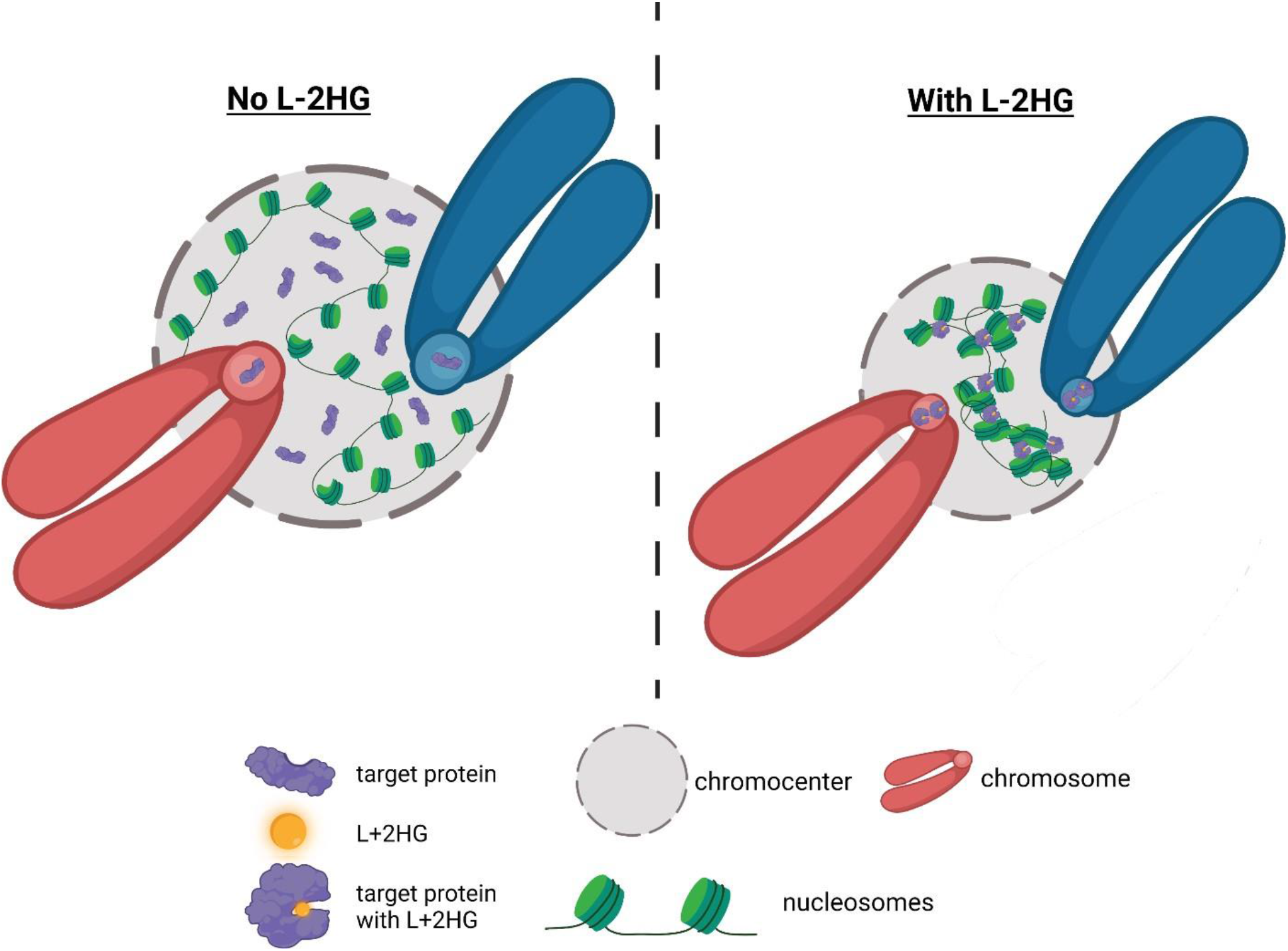
Hypothetical model for the effects of L-2HG on chromocenters and centromeres. upon binding of L-2HG the target protein undergoes a conformational change. Some of the L-2HG bound target protein translocates from the chromocenter to the centromere. The L-2HG bound target induces condensation of the nearby chromatin. This could occur either via protein-protein interactions or via induction of polymer-polymer phase separation as was shown previously for HP1α. This figure was Created with BioRender.com.

## Acknowledgments

We thank Prof. Norman Grover for help with some of the statistical analyses and Drs Ittai Ben-Porath, Yuval Dor, Yotam Drier and Yehudit Bergman for helpful discussions and comments on the manuscript. We thank Mohammad Jumaa for excellent histological assistance, Dr. Douglas Lutz, Getter Group BioMed and Dr. Abel Pereira da Graça, Carl Zeiss Microscopy, GmbH, for help with high resolution microscopy analyses, Dr. William Breuer for his assistance in proteomics analysis and Dr. Yuval Nevo for assistance with bioinformatic analyses.

## Author Contributions

Conceptualization: NM, MS, TS, MK, EP; Methodology: NM, YM; Statistical and bioinformatics analyses: NM, NP; Investigation: NM, MS, YM, IS, FM, GB, EM, ZM, EA, BS, JM. Writing original draft: NM, EP; Writing - Review & Editing: NM, MS, GB, NP, IS, TS, MK, EP. Visualization: NM, MS, YM, NP. Supervision: NM, TS, MK, EP; Funding acquisition: EP.

## Materials and Methods

### Methods

#### Mice

Stra8-icre (B6.FVB-Tg (Stra8-icre)1Reb/LguJ) male mice (stock 017490), tdTomato (B6;129S6-Gt(ROSA)26Sortm9(CAG-tdTomato)Hze/J females(stock 007905) and EYFP (B6.129X1-Gt(ROSA)26Sortm1(EYFP)Cos/J females (stock 006148) were purchased from Jackson Laboratories. All mice are on C57BL/6 background. Hemizygous Stra-icre males were mated with either homozygous tdTomato or with homozygous EYFP females, and males which were positive for Stra-icre were used as a source for testis tissue. Genotyping was done by quantitative PCR according to the suppliers’ protocol. (https://www2.jax.org/protocolsdb/f?p=116:5:0::NO:5:P5_MASTER_PROTOCOL_ID,P5_JRS_CODE:29550,017490). When indicated, mice were treated with 1.3 g/kg oxamate i.p. once daily, for the indicated periods of time. All animal studies were approved by the Hebrew University Institutional Animal Care and Use Committee.

#### Cell Isolation

Prepubertal and adult (above 10 weeks old) mice were euthanized with pental, and cervical dislocation was performed. Testis were removed, put into cold Hank’s solution and seminiferous tubules were released from the tunica albuginea. Tubules were examined with fluorescence microscope to verify tomato expression. Tissue digestion was performed in two steps. First, tubules from two testes were put into 20 ml medium (DMEM/F12) with collagenase (1 mg/ml) and DNAse (0.28 mg/ml), then gently dispersed by pipetting using sequentially 25, 10 and 5 ml pipettes. After incubation for 2 min in room temperature and washing with medium, 30 ml of trypsin diluted 2:1 with medium and containing 0.2 mg/ml DNAse was added. Tubules were incubated at 32°C for 5 min with frequent gentle inversions of the tube. After increasing of DNAse concentration to 0.93 mg/ml and additional 3 min incubation in room temperature, 2.8 ml FBS was added to stop trypsin activity. The resulting suspension was passed through 40 μm cell strainer, washed in medium and incubated with dead cells removal beads for 15 min in room temperature. Live cells were collected according to the manufacturer’s instructions and washed in PBS with 5% serum followed by dispersal in sorting buffer. When separation between Undiff and Diff Spg was required, live cells were pre-incubated for 10 min on ice with 1% rat serum and 1.2% FC in PBS, to block non-specific binding, continued by incubation with APC-cKit antibody (1: 900) for 20 min on ice.

#### Flow Cytometry Sorting

After live cell isolation, cells were washed in 2% dialyzed serum in PBS, resuspended in “sorting buffer” at a density of 6 million cells / 1 ml. During sorting, cells were collected into 5 ml tubes, which were kept overnight in 4°C, inverted with 2ml of different “collecting buffers” to prevent cell loss. Cells were sorted using a BD FACSAria™ III sorter. Debris exclusion was done based on light scattering. Tomato intensity (yellow-green laser, bandpass582/15) of cells versus forward scatter (FSC) was used to differentiate between the different populations. Doublets exclusion in each population was performed on the basis of FSC width/area (FWA), followed by side scatter (SSC) width/area (SWA) and tomato width/area (PWA). APC-cKit positive cells were detected using red laser, bandpass 660/20. Ckit-negative gates were established using a sample of cells incubated with APC-Rat IgG instead of APC-cKit. Four-way sorting was performed using 100-micron nozzle and threshold was kept in the range of 2500-3000 events/second as PDD cells were fragile and sensitive to high pressures that can develop during sorting. Characterization and purity assessment of sorted cells was done by measurement of DNA content of the isolated populations (before and after sorting) and staining for specific markers of whole cells and nuclear spreads, as described below. Purity of populations was calculated by counting at least 100 stained cells or nuclear spreads using ImageJ software and by DNA content estimated by propidium iodide after sorting. The PDD population consists of about 6% diakinesis cells. For studies of the diakinesis stage we isolated cells that appeared in high FSC area at the gate of PDD. We achieved about 50% enrichment of these rare cells.

#### DNA content

During sorting, DNA content estimation was performed by incubating cells (directly after dead cell removal procedure) with 10 μg/ml Hoechst for 30 min, 32°C in the dark. In sorted and isolated populations, DNA content was estimated by PI staining. Cells were washed with cold PBS and dispersed in 250 μl of PBS. Cells were fixed by adding 0.75 ml of 100% high purity ethanol dropwise while slowly vortexing to a final volume of 75% ethanol. Fixed cells were washed twice with 1 ml cold PBS and spun down at 500g for 10 min at 4°C, after each wash. Cells were resuspended in 0.2 ml “PI-Mix” (**Materials**) and incubated for 15-30 min in the dark before analysis. tdTomato fluorescence disappears after ethanol fixation, allowing us to detect PI wavelength in tdTomato/PE channel (**Figure S1D**).

#### Immunofluorescence staining of sorted cells for identification of early meiotic populations

Isolated cells were attached to poly L-lysine pretreated slides (1 mg/ml, 10 min) using cytospin (500 rpm, 3 min, low velocity to prevent PD cell damage). Cells were fixed by dipping the slides in methanol at-20°C for 30 sec. Slides were washed three times with PBS (3 min), then washed for 10 min in 0.1% triton X-100 in PBS and blocked with 3% BSA, 5% donkey serum and 0.1% triton X-100 in PBS for 1 hr. Primary antibody solution was prepared in 0.1% triton X-100, 1%BSA in PBS, rabbit anti PLZF 1:100, mouse anti DMRT1 1:100, and slides were incubated overnight at 4°C. After overnight incubation, slides were washed 3 times for 10 min in PBS 0.1% triton X-100 and incubated with secondary antibodies: FITC conjugated anti-mouse 1:1000 and Cy5 conjugated anti-rabbit 1:500 in PBS with 1% BSA for 2 hr in dark at room temperature. Following 3 washings for 3 min in PBS, slides were covered with mounting solution containing DAPI and imaged with an Olympus FV3000 confocal microscope. The prevalence of PLZF and DMRT1 expressing cells was evaluated after counting at least 100 cells using ImageJ (**Figure S1E-H**).

#### Nuclear spreads preparation and immunofluorescence staining

Nuclear spreads were prepared as described^29^ with modifications as follows: Cells were collected into Eppendorf tubes and washed once with PBS (350g, 5min) and all liquid was removed. Cells were dispersed in 150μl of “hypotonic solution” (**Materials**) containing 0.5 mM PMSF, protease inhibitor cocktail X1 and 0.5 mM DTT, then 150μl of PBS was added and tubes were incubated in rt for 8 min with periodic flickering. After incubation, 1.2 ml PBS was added and cells were spun for 5min 1000g. Most of the liquid was removed, 8μl of the left liquid was moved to a new Eppendorf tube and 16μl of sucrose 100mM, pH=8.2 was added. 10-20 thousand nuclei in 6μl of the above solution were spread dropwise on a framed 2X1.5 cm area on a slide covered with 42μl freshly prepared and filtered solution of 1% PFA, 0.15% Triton X-100, 1 mM sodium borate, pH= 9.2. Slides were put in closed humid chambers overnight. Next day, slides were air dried for approximately 1hr, washed twice in 0.4% Photo-flo in water (2min), then washed 3 times in water (1 min) and stored in PBS 4°C until staining.For immunofluorescence staining, slides were washed for 5 min in 0.5% triton X-100 in PBS, 5 min in 0.1% triton X-100 in PBS and blocked in filtrated 2% BSA, 0.1% triton X-100, 10% donkey serum in PBS for 1hr. Primary antibodies were prepared in filtrated 2% BSA, 0.1% triton X-100 and 10% donkey serum: rabbit anti-SCP3 1:750, mouse anti-γH2AX 1:3000, rabbit H3K9me3 1:500, rabbit HP1α 1:250, human CREST serum 1:20, rabbit LDHC 1:50, rabbit LDHA 1:100, then slides were incubated overnight in 4°C. Next day, slides were washed three times for 5 min in 0.1% triton X-100 in PBS, 5 min in PBS. Secondary antibodies were diluted in PBS with 1% BSA: FITC conjugated anti-mouse 1:500, CY5 conjugated anti-rabbit 1:500, Alexa 647 conjugated anti-human 1:100 and slides were incubated for 2 hr in dark at rt then washed three times for 5 min PBS, followed with mounting solution containing Hoechst 1:250 (Vector) or Prolong Glass antifade with NucBlue. After staining, chromosomes were observed under Nikon Eclipse Ti microscope at X100 (**Figures: 5A, 6A, S1E-H and S3A,D**) or LSM 980 Zeiss Airyscan2 at X63 magnification (**Figures: 4B-J, 5C-F, S2C, S3B and S4A-D**). Evaluation of stained area and its intensity was performed using NIS elements or Zeiss 980 analyzer software. Identification of pachytene and diplotene cells was done by evaluation of the presence of synapsed and asynapsed regions of the bivalents based on SCP3 staining. In pachytene stage bivalents are fully synapsed, while in diplotene, bivalents are partially asynapsed.

#### Fresh Cell Nuclear Spreads

Cells were isolated as described above and after dead cells removal adhered to slides using cytospin and fixed by dipping in methanol at −20°C for 10 sec. Using this procedure chromosomes could be visualized in whole cells. In addition, some PD cells released chromosomes, which could be analyzed using different antibodies for staining. Slides were washed for 5 min in PBS and blocked in filtrated 3% BSA, 0.1% triton X-100, 5% donkey serum in PBS for 1hr. Primary antibodies were diluted in blocking solution: mouse anti-SCP3 1:200, LDHC 1:50, LDHA 1:100, slides were incubated overnight in 4°C with antibodies. Next day, slides were washed three times for 10 min in 0.1% triton X-100 in PBS, 5 min in PBS. Secondary antibodies were diluted in PBS with 1% BSA: FITC conjugated anti-mouse 1:500, CY5 conjugated anti-rabbit or anti-goat 1:500, and slides were incubated for 2 hr in dark at rt then washed three times for 5 min in PBS. Mounting was performed using ProLong Glass Antifade and after 48 hrs curation pictures were taken with LSM 980 Zeiss Airyscan confocal microscope at X 63 magnification (**Figures: 3A,C and S2B,D**). Stimulated Emission Depletion Microscopy (STED) samples were prepared as described above, using Abberior STAR RED and Abberior STAR ORANGE secondary antibodies. Images were taken with an Abberior Instruments Facility LINE microscope equipped with an inverted IX83 microscope (Olympus), a 60x oil objective (UPlanXApo 60x/1.42 oil, Olympus), using pulsed excitation lasers at 561 nm and 640 nm and a pulsed STED laser operating at 775 nme. All acquisition operations were controlled by Lightbox Software. STED images presented in the figures were only adjusted in brightness and contrast on raw data using Fiji software 6 (**Figures: 3B, D and 5B**).

#### Histological sections

Testes were removed, fixed in formalin (for immunofluorescence and immunohistochemistry) or Bouin fixative (for hematoxylin eosin, PAS or Caspase 3 stains) overnight and embedded in paraffin (**Figure S5**). Histological sections were stained with mouse anti histone H3 phosphoserine 10 (PHH3) 1:400 and rabbit anti MAD1 1:100 after antigen retrieval in citrate buffered and blocking in filtrated 2% BSA, 0.1% triton X-100, 10% donkey serum in PBS for 1hr (**Figures 6C,E**).

#### Culturing of isolated cell populations

LZ and PD cells were cultured in “culturing medium” for LZ and PDD (**Materials**), at 32°C as described^53^. L –or D-2HG-octyl, final concentration 0.3 mM or 0.6 mM, for several hours or 10 min incubations, respectively, was added dissolved in DMSO, 1.5 μl/1 ml medium, sodium oxamate, final concentration 24 mM, was added dissolved in water, 54 μl/1 ml medium. Untreated cells received equivalent volumes of solvents. When LZ cells were cultured for more than 48 hours, half of the medium was replaced with fresh medium containing drug or vehicle every two days.

#### Gene expression-RNAseq

Total RNA of each sorted population from three independent experiments was isolated using Quiagen mini kit according to the manufacturer’s instructions. PolyA mRNA was captured using magnetic oligo dT beads followed by fragmentation and cDNA synthesis. Libraries were constructed from fragmented DNA while performing end repair, A-base addition, adapter ligation and PCR amplification steps with SPRI beads cleanup in between steps. Indexed samples were pooled and sequenced in an Illumina HiSeq 2500 machine in a single read mode. Median sequencing depth was ~18 million reads per sample. Sequencing depth was homogenous across samples in the experiment. Performed by G-INCPM (Weizmann Institute of Science).

#### Tracing with UC13-L-lactate, UC13-D-glucose and UC13-glutamine

Cells were sorted into PBS with 5% dialyzed serum. LZ and PDD cells (between 0.5 to 1 million cells) were incubated in 200 μl of K salt solution supplemented with 10 % dialyzed serum, 2mM glutamine and with 5 mM UC13 L-lactate, UC13 D-glucose or 2 mM UC13 glutamine in Eppendorf tubes for 120 min, at 32°C. At the end of the incubation cells were washed twice in PBS, and extracted in methanol-acetonitrile 5:3. Extracted metabolites were separated on a SeQuant ZIC-pHILIC column (2.1 × 150 mm, 5 μm bead size, Merck Millipore). Flow rate was set to 0.2 mL/min, column compartment temperature was set to 30°C, and autosampler tray was maintained at 4°C. Mobile phase A consisted of 20 mM ammonium carbonate with 0.01% (v/v) ammonium hydroxide. Mobile Phase B was 100% acetonitrile. The mobile phase linear gradient (%B) was as follows: 0 min 80%, 15 min 20%, 15.1 min 80%, and 23 min 80%. A mobile phase was introduced to Thermo Q-Exactive mass spectrometer with an electrospray ionization source working in polarity switching mode. Metabolites were analyzed using full-scan method in the range 72-1,080 m/z and with a resolution of 70,000. Ionization source parameters were as following: sheath gas 25 units, auxiliary gas 3 units, spray voltage 3.3 and 3.8 kV in negative and positive ionization mode respectively, capillary temperature 325 °C, S-lens RF level 65, auxiliary gas temperature 200 °C. Positions of metabolites in the chromatogram were identified by corresponding pure chemical standards. Data was analyzed using the MAVEN software suite^54^. Relative metabolite levels were quantified from peak areas and normalized to unit mean for each peak after correction for protein content (**Figures 2A-B and 4A).**

#### Identification of 2HG enantiomers

Cell extracts were prepared using 50%MeOH+30%AcN+20%PBS mixture. Evaporated standards and samples were derivatized with TSPC by chiral derivatization approach and evaporated. For analysis, samples were re-dissolved in 100μL of 30%-aqueous acetonitrile containing phthalic acid as internal standard (10μM), centrifuged twice at 21,000*g* for 5 min to remove insoluble material. The LC–MS/MS instrument consisted of an Acquity I-class UPLC system (Waters) and Xevo TQ-S triple quadrupole mass spectrometer (Waters) equipped with an electrospray ion source and operated in positive ion mode. MassLynx and TargetLynx software (version 4.1, Waters) were applied for data acquisition and analysis. Chromatographic separation was done on a 100 mm × 2.1 mm internal diameter, 1.7-μm UPLC BEH C8 column equipped with 50 mm × 2.1 mm internal diameter, 1.7-μm UPLC BEH C8 pre-column (both Waters Acquity) with mobile phases A (0.1% formic acid) and B (0.1% formic acid in 95%-aqueous acetonitrile) at a flow rate of 0.25 ml min-1 and column temperature 25°C. A gradient was used as follows: 0–1 min a linear increase from 30 to 32% B, then held at 32% B till 9.5 min, increase to 80% B during 1.5 min, then back to 30% B in 0.5 min, and equilibration at 30% B for 2.5 min, providing total run time of 13 min. Samples kept at 8 °C were automatically injected in a volume of 3 μl. MS parameters (negative polarity): capillary voltage - 2.4kV, source temperature - 120°C. MRM transitions (collision energy, eV): 448.0>155.0 (29) and 448.0>318.0 (13) for derivatized 2HG, 446.0>155.0 (29) and 165.0>77.0 (15), 165.0>121.0 (10) for internal standards. Performed by Life Science Core Facilities, Weizmann Institute of Science (**Figure S2A).**

#### Cell volume measurements

Cell diameters were measured by two independent methods using either microscopy or an Eclipse FACS analyzer. Images of the isolated populations were taken using Nikon Eclipse Ti microscope at X400 magnification. Diameters of 100 cells were estimated using ImageJ tool. Mean area and volume were calculated from the average diameter of cells in each population. In addition, all cells sizes (diameter, area and volume) were measured by an Eclipse FACS analyzer. The analyzer was calibrated using a Flow Cytometry Size Calibration Kit. We used the average of the diameters estimated by these two methods to calculate cell area and volume (**Table S1**).

#### Quantitative PCR of minor and major satellite transcripts

Cells were harvested with Trizol for RNA extraction. RNA was prepared using miRNeasy mini kit (Qiagen-cat no 217004). Reverse transcription was carried out using SuperScript III First-strand Synthesis System (Invitrogen). The quantification of PCR products was analyzed with SYBRgreen using ABI PRISM 7900 Sequence Detection system software (Applied Biosystems). The primer sequences were MajSAT 5’-GGCGAGAAAACTGAAAATCACG-3’,5’-CTTGCCATATTCCACGTCCT-3’; MinSAT5’-TGGAAACGGGATTTGTAGA-3’, 5’-CGGTTTCCAACATATGTGTTTT-3’. Results were normalized to Tox4 5’-TCCCGGAGGAAATGACAATTACC-3’, 5’-GTGAGGGATCAGAGTCCAAGG-3’ and Brd2 5’-AATGGCTTCTGTACCAGCTTTAC-3’, 5’-CTGGCTTTTTGGGATTGGACA-3’^50^ (**Figures: 5I-J and S4E-F**).

#### HP1α and histone modifications

PDD cells were isolated and incubated for 48 hours with vehicle, 24 mM oxamate or oxamate with 0.3 mM L-2HG in duplicates. Cells were lysed in RIPA containing 1X protease inhibitors (Roche). Lysates were frozen in −20°C. Following de-freezing they were sonicated for 3X30 sec and cleared by centrifugation at 14,000 x g for 15 min at 4°C. An equal amount of protein was extracted from 400K cells and loaded onto a 12% acrylamide gel. HP1 and SCP3 were probed with their respective antibodies, SCP3 was used to normalize HP1α ammounts, (**Figure 5G-H**). Histone lysine modifications were measured using a histone extraction kit ab113476 and multiplex colorimetric assay kit ab 185910, 25 ng protein/well (**Figure S4E**).

#### Sample preparation for the whole cell proteomic mass spectrometry (MS) analysis

5-6 replicates of 300 000 LZ and 150 000 PDD cells were isolated from 5 mice and collected via sorting into PBS buffer. Cells were lysed with 5% SDS, 50 mM Tris-HCl pH 8.0, sonicated and centrifuged at 14000 X g for 15 min in room temperature (**Figure 2D**).

#### Isolation of chromatin-bound proteins for proteomic MS analysis

The isolation of chromatin –bound proteins was performed with the modification of a method described by^27,28,55^. Composition of the solutions used is listed in “Materials”. 6 X10^6^ LZ cells and 12 X 10^6^ PDD cells were isolated from 6 mice. Cells were homogenized in homogenization buffer followed by a 1000 X g centrifugation for 5 min to isolate nuclei. Low salt extraction (LSE) was performed by vigorous mixing nuclei in 1 ml LSE-buffer and an 100 000 X g, 4 deg centrifugation for 30 min. The supernatant was discarded and the precipitate containing chromatin was washed twice with 1 ml washing buffer (WB), mixing by vortex and centrifugation as described above. Chromatin bound proteins were extracted with high salt extraction (HSE) buffer added by extensive sonication. After centrifugation, as described above, HSE was collected for MS.

(**Figure 3E**).

#### LC MS/MS analysis

The soluble extracts were supplemented with 10 mM dithiothreitol and incubated for 10 min. at 80°C. The proteins were alkylated by addition of 55 mM iodoacetamide (Sigma Chem. Corp. St. Louis, MO) and incubation for 30 min. at room temperature in the dark. Removal of SDS followed by digestion with sequencing grade modified trypsin (Promega Corp., Madison, WS) were performed using the S-Trap microspin column kit as specified by the manufacturer (Protifi,LLC, Huntington, NY). The tryptic peptides were desalted on C18 Stage tips (Rappsilber et al., 2007). A total of 0.3 μg of peptides (determined by Absorbance at 280 nm) from each sample were injected into the mass spectrometer.

MS analysis was performed using a Q Exactive Plus mass spectrometer (Thermo Fisher Scientific) coupled on-line to a nanoflow UHPLC instrument (Ultimate 3000 Dionex, Thermo Fisher Scientific). Eluted peptides were separated over a 120-min gradient run at a flow rate of 0.15 μl/min (during the separation phase) on a reverse phase 25-cm, C18 column (75μm ID, 2μm, 100Å, Thermo PepMap®RSLC from Thermo Scientific). The survey scans (380–2,000 m/z, target value 3E6 charges, maximum ion injection times 50 ms) were acquired and followed by higher energy collisional dissociation (HCD) based fragmentation (normalized collision energy 25). A resolution of 70,000 was used for survey scans and up to 15 dynamically chosen most abundant precursor ions were fragmented (isolation window 1.8 m/z). The MS/MS scans were acquired at a resolution of 17,500 (target value 5E4 charges, maximum ion injection times 57 ms). Dynamic exclusion was 60 sec.

Mass spectra data were processed using the MaxQuant computational platform, version 1.6.17.0. Peak lists were searched against the mouse Uniprot FASTA sequence database of reviewed entries from Mar. 4, 2021, containing 36,759 entries. The search included cysteine carbamidomethylation as a fixed modification and oxidation of methionine and N-terminal acetylation as variable modifications. Peptides with minimum of seven amino-acid length were considered and the required FDR was set to 1% at the peptide and protein level. Protein identification required at least 2 unique or razor peptides per protein group. Relative protein quantification in MaxQuant was performed using the label free quantification (LFQ) algorithm^56^. LFQ in MaxQuant uses only common peptides for pair-wise ratio determination for each protein and calculates a median ratio to protect against outliers. It then determines all pair-wise protein ratios and requires a minimal number of two peptide ratios for a given protein ratio to be considered valid.

Statistical analysis (*n=6*) was performed using the Perseus statistical package^57^. The Perseus computational platform for comprehensive analysis of (prote)omics data. Only those proteins for which at least 3 valid LFQ values were obtained in at least one sample group were accepted for statistical analysis by Volcano plot (t-test, *p* < 0.05). After application of this filter, a random value was substituted for proteins for which LFQ could not be determined (“Imputation” function of Perseus). The imputed values were in the range of 10% of the median value of all the proteins in the sample and allowed calculation of p-values, (**Figures 2D-E and 3E).**

#### Statistical analysis

All experiments were repeated independently at least twice with similar results. For normal variable evaluations, data was shown as mean value of replicates with their respective standard errors (mean±SE); for non-parametric variable evaluations, data was shown as median with range as indicated in the figure legends. Student’s t test, Mann Whitney, Friedman, Bonferroni and Tukey tests were performed as appropriate. P values were calculated by using Graphpad Prism 9 software (San Diego, CA, USA). Levels of significance are indicated as *p < 0.05; **p < 0.01; ***p < 0.001; ****p < 0.0001. Statistical details for all experiments can be found in the respective figure legends. All tests were two tailed unless indicated otherwise. No statistical methods were used to predetermine sample size. Unless otherwise stated, experiments were not randomized and investigators were not blinded to allocation during experiments and outcome assessment.

### Materials

(2S)-Octyl-α-hydroxyglutarate (HY-103641) MedChemExpress dissolved in DMSO

(R)-Octyl-α-hydroxyglutarate (SML2200) Merck dissolved in DMSO

Bovine serum albumin fatty acids free (A8806-5G) Sigma

Clarity Western ECL Substrate (170-5061) Bio-Rad

Collagenase (C5138) Sigma dissolved in DMEM/F12 medium 8 mg/ml

Collecting Buffer (during sorting) for experiments in which cells were cultured consisted of DMEM/F12 supplemented with 10% dialyzed serum, 3mM L-lactate, 3mM pyruvate and antibiotic-antimycotic solution.

Collecting Buffer (during sorting) for gene expression consisted of PBS with 2% albumin Collecting Buffer (during sorting) for tracing studies consisted of PBS with 5% dialyzed FBS. Culturing medium (LZ and PD): MEM eagle (Ref # 01-040-1A from BI) supplemented with 2 mM glutamine, 5mM sodium lactate, 5mM sodium pyruvate, 15mM HEPES, 1% antibiotic-antimycotic (Thermo Scientific, 15240096) v/v, 5% dialyzed serum.

Dead cells removal kit (130-090-101) Miltenyi

Dialyzed Serum (040111B) Biological Industries

DNAse (DN-25) Sigma dissolved in saline 7 mg/ml FBS

heat inactivated (04-127-1) Biological Industries

Flow Cytometry Size Calibration Kit (F13838) Thermo Fisher Scientific Gels

for Western 4568094 Bio-Rad

Hoechst 33342 (H1399) Invitrogen, stock solution 1mg/ml.

Solutions for extraction of chromatin-bound proteins for proteomic mass spectrometry analysis:

1. Homogenization solution - 0.32 M sucrose, protease inhibitors cocktail, 1 mM PMSF and 10 mM HEPES, pH 7.4.
2. Low salt extraction buffer (LSE)- 100 mM KCl, 0.4 mM EDTA, 0.1% Triton X-100, 10% glycerol,1 mM mercaptoethanol, protease inhibitors cocktail and 20 mM TrisHCL, pH 7.5
3. Wash buffer (WB)- 1mM CaCl2, 1.5 mM MgCl2, 10 % glycerol and 10 mM TrisHCl, pH 7.5.
4. High salt extraction (HSE)- 400 mM KCL, 5 mM MgCl, 2% Tween-20, 10% glycerol, 1mM mercaptoethanol, protease inhibitors cocktail and 20 mM Hepes-KOH, pH 7.

Hypotonic solution (for spreads): 30mM tris-HCl, 100 mM sucrose, 17mM sodium citrate, 5mM EDTA, PH=8.3

K-salt solution: 142 mM NaCl, 5.4 mM KCl, 0.91 mM NaH2PO4, 1.8 mM CaCl2, 0.8 mM MgSO4 and HEPES 5mM, pH 7.35 (*P/D cells are sensitive to Mg concentration. Higher than 0.8 mM Mg can cause high death rate)

Medium DMEM/F12 01-170-1A Biological Industries miRNeasy mini kit (217004) Qiagen

Mounting solution for immunofluorescent staining ProLong Glass Antifade P36983 Invitrogen

Photo-flo (501 0640) KODAK

PI-mix consists of PBS with (50μg/ml) Propidium Iodide and 50μg/ml RNAse-A Poly

L-lysine (p4707) Sigma.

Propidium Iodide- PI (P1304MP) Invitrogen, stock solution 250 μg/ml.

Protein assay- (5000006) Bio-Rad

Rat serum (24-5555-4) and FC (14-0161-82) eBioscience

RNAse-A (9001-99-4) Sigma

RNeasy micro kit (74004) Qiagen

RNeasy mini kit (74104) Qiagen

Sodium L-lactate 13-C3 (CLM-1577-0.5) Cambridge isotopes laboratories.

Sodium oxamate (O2751) Sigma

Sorting buffer consisted of PBS with 25 mM HEPES, 5% dialyzed FBS, 0.05 mg DNAse and 0.75 mM MgCl2

Trans-Blot Turbo Transfer Pack-(1704157) Bio-Rad

Trypsin-EDTA (03-0521-1) Biological Industries

| antibody | Cat # |
| --- | --- |
| Caspase 3 | 9661 (Cell Signaling) |
| CD117 (cKit- APC conjugated) | 105812 (Biolegend) |
| CENP-A | C51A7 (Cell Signaling) |
| CENP-C | NBP2-75438 (Novus) |
| CENP-N | 16751-1-AP ((Proteintech) |
| Crest | 15235 (Antibody INC) |
| DMRT1 | SC-377167 (Santa Cruz) |
| H3K9me3 | ab8898 (Abcam) |
| HP1α | AB-ab109028 (Abcam) |
| LDHA | 199871-A (Proteintech) |
| LDHC | 199891-A (Proteintech) |
| LDHC | ab3966 (Abcam) |
| PhosphoH3 (serine10) | 36916 (Cell Marque) |
| PhosphoH3 (serine 10) | MA5-15220 (Invitrogen) |
| PLZF | AB5535 (Millipore) |
| SCP3 | ab15093 (Abcam) |
| SCP3 | ab97672 (Abcam) |
| SOX9 | AB5535 (Millipore) |
| γH2AX | 05-636 (Millipore) |
| MAD1 | 18322-1-AP (Proteintech) |
| STAR ORANGE | 00206JR-3 (Abberior) |
| STAR RED | 00121JR-1 (Abberior) |
| Cy5 donkey anti rabbit | ab150075 (Abcam) |
| Alexa488 goat anti mouse | Ab150117(Abcam) |
| Cy5 donkey anti human | 709175 149 (Jackson) |

