## Supplementary figures for "2-hydroxyglutarate controls centromere and heterochromatin conformation and function in the male germline"

Supplementary figures: **2-hydroxyglutarate controls centromere condensation and function in the male germline.**

Mayorek et al.
2022

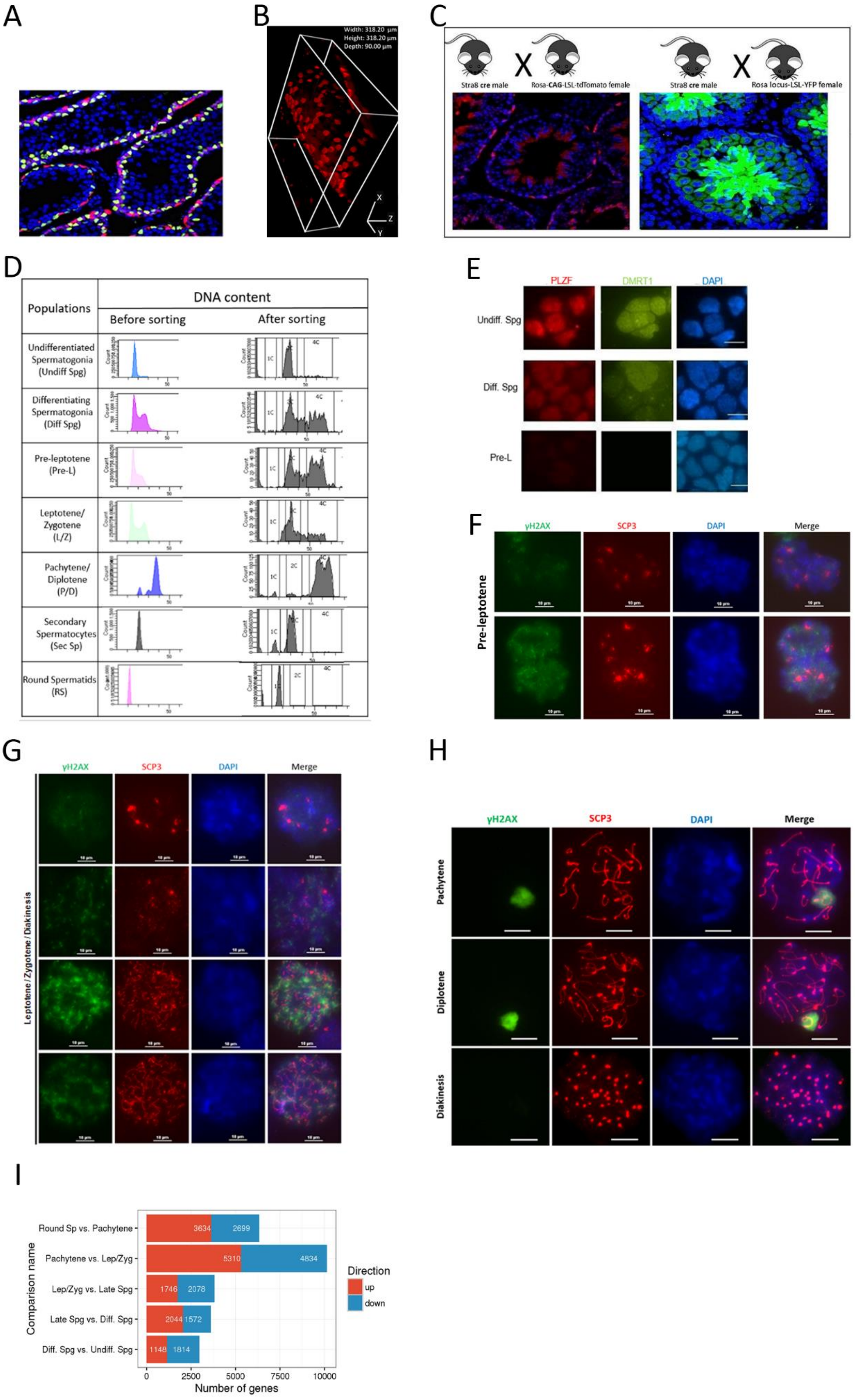

**Figure S1: Stra8-Tom mice allow isolation of highly purified populations of germ cells along their differentiation process. A**. tdTomato fluorescence (red) and immunostaining for Sox9 (green), demonstrating germ cells specificity of tdTomato expression, frozen section of Stra8-Tom mouse. **B.** 3D reconstruction of a whole mount preparation of seminiferous tubule, demonstrating decreasing tomato fluorescence towards the lumen. **C.** Stra8-Tom mice were produced by mating Stra8-icre males with CAG-LSL-tdTomato females (left). Stra8 icre-RosaYFP males were produced by mating Stra8 icre males with Rosa-LSL-YFP females (right). Shown are frozen sections counterstained with DAPI. Note that while the Rosa locus drives expression throughout the sperm lineage, expression driven by the CAG promoter gradually diminishes. The luminal fluorescence represents autofluorescence of sperm tails. **D.** DNA content was measured by adding Hoechst to testis cell suspensions followed with FACS analysis based on tdTomato intensity (before sorting) vs. Hoechst. Alternatively, cell populations were sorted as described, ethanol- fixed and stained with propidium iodide. The purity of cell populations based on DNA content after sorting was estimated as following: Undiff.Spg-100% 2C, Diff.Spg, PreL and LZ – mixed population of cells between 2C (incomplete replication) and 4C (complete replication), PDD – mostly 4C cells with slight contamination with 2C and about 14% 1C, Sec.Sp-85% 2C and 15% 1C, RS-100% 1C. **E.** Isolated populations were attached to slides by cytospin centrifugation and immunostained for PLZF and DMRT1 (markers of spermatogonia). 90% of undifferentiated spermatogonia (ckit negative) were strongly positive for both markers, 90% of differentiating spermatogonia (ckit positive) were weakly positive for both markers and absent in all other populations. Changes between undifferentiated and differentiating spermatogonia (separated using ckit antibody) were corroborated by qPCR (data not shown). PLZF expression was 4.4, 4.8 and 10 fold higher in Undiff.Spg versus Diff.Spg in 3 respective experiments. GFR-α (another Undiff.Spg marker) expression was 8.7, 10 and 14-fold higher in Undiff.Spg. **F.** PreL cells were isolated and nuclear spreads were stained with antibodies for SCP3 and γH2AX. γH2AX staining was absent (upper panel) or present (lower panel). **G.** LZ cells were isolated and nuclear spreads were stained with antibodies for SCP3 and γH2AX. 44% of cells do not express γH2AX (PreL kind of cells, first row). 56% are a mixture of cells expressing different levels of γH2AX and different levels of synaptonemal complex formation (estimated by counting of 100 cells). **H.** PDD cells were isolated and nuclear spreads were stained with antibodies for SCP3 and γH2AX. The average composition of this population is: 40% pachytene, 40% diplotene, 6% diakinesis and 14% round spermatids. Scale bar **E-H**: 10 µm**. I**. mRNA was extracted from the freshly isolated indicated populations of 3 mice. Shown are the numbers of genes which are differentially expressed (at least 2-fold difference, FDR<0.05) between the indicated populations**.**

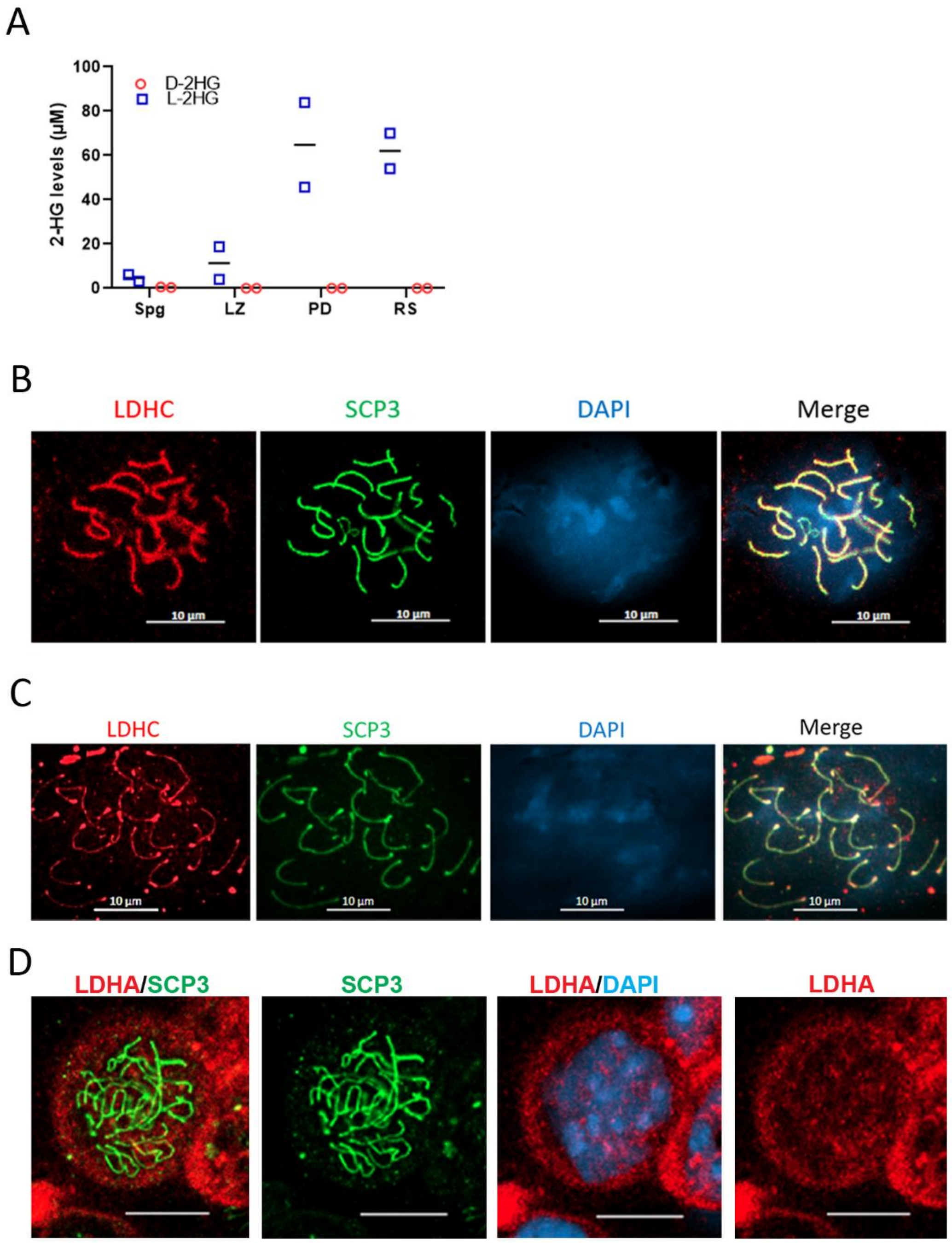


**Figure S2: LDHC generates L-2HG, is expressed in PDD cells and can be found in the nucleus. A.** L- and D-2HG enantiomer content was measured in the indicated populations using chiral derivatization. Results of two independent experiments are shown. Calculation of 2HG concentrations was performed using the volumes of cells in **Table S1**. D-2HG enantiomer was detected only in Spg cells at negligible concentrations of 0.6 and 0.4 µM. **B-F**: Verification of LDHC staining along chromosomes and in centromeres. cells. **B.** Freshly isolated testicular cells (the same procedure as shown in **Figure 3**), LDHC stained with goat ab3966 antibody (and not rabbit Proteintech 19981-A) and co-stained with SCP3. **C.** Spreads prepared using Papanikos et al., protocol described in methods [29]. LDHC was stained with rabbit Proteintech 19981-A and co-stained with SCP3. Strong staining of centromeres with LDHC can be noted. Scale bar 10 µm. **D:** LDHA is not localized on chromosomes.Confocal microscope image of a diplotene cell stained with LDHA antibody, scale bar 10 µm, compare with **Figure 3A**.

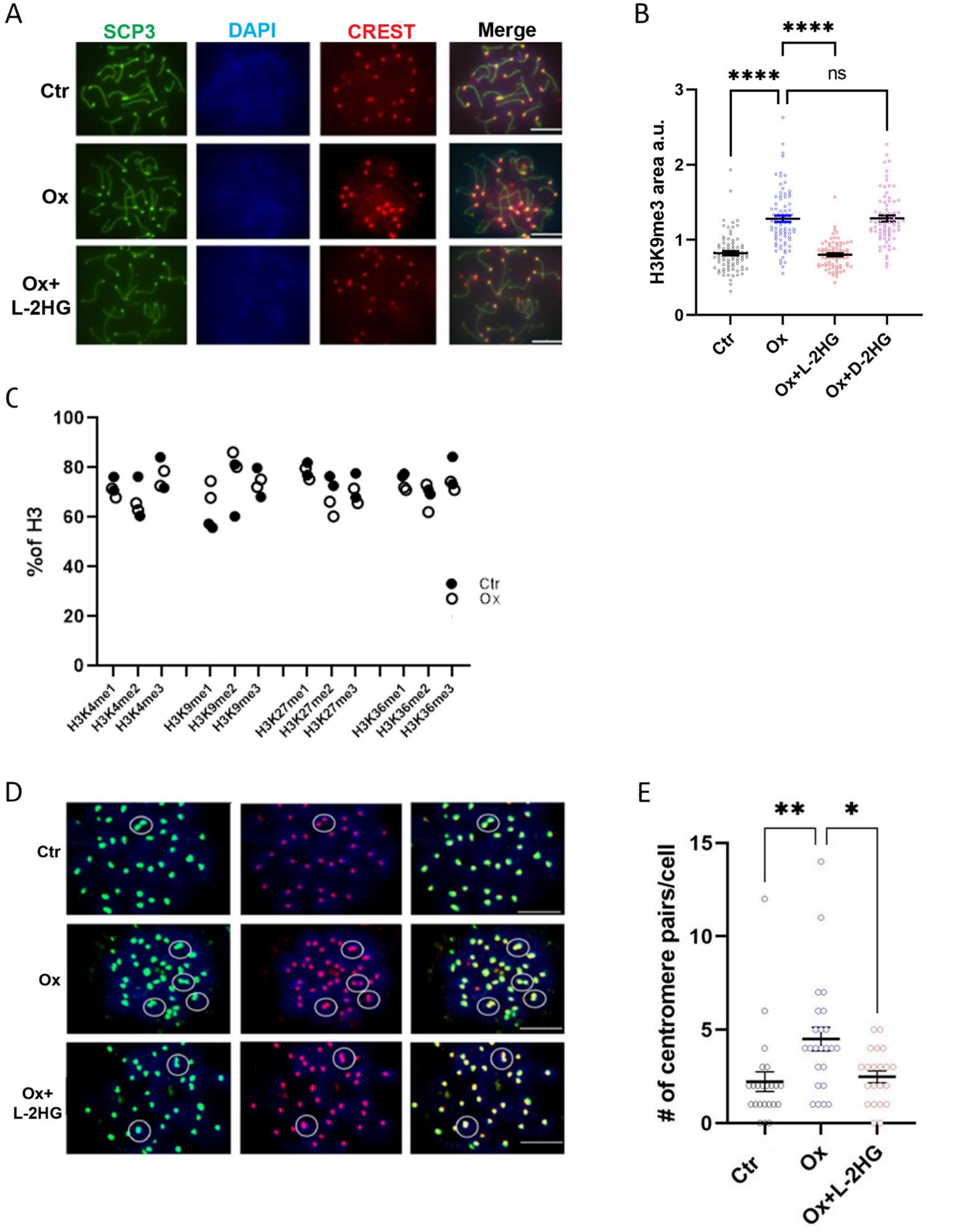

A
SCP3
DAPI
CREST
Merge
Ctr
Ox
Ox+
L-2HG
B
****
****
ns
H3K9me3 area a.u.
0
1
2
3
Ctr
Ox
Ox+L-2HG
Ox+D-2HG
C
%of H3
0
20
40
60
80
100
Ctr
Ox
H3K4me1
H3K4me2
H3K4me3
H3K9me1
H3K9me2
H3K9me3
H3K27me1
H3K27me2
H3K27me3
H3K36me1
H3K36me2
H3K36me3
D
Ctr
Ox
Ox+
L-2HG
E
**
*
### of centromere pairs/cell
0
5
10
15
Ctr
Ox
Ox+L-2HG

**Figure S3: L-2HG is necessary for accurate centromere and chromocenter morphology in PDD cells.** Isolated PDD cells were incubated for 24 (B and D-E) or 48 hours (A and C) with vehicle, 24 mM oxamate or 24 mM oxamate with 0.3 mM cell-permeable L-2HG, as indicated. **A**. Representative confocal microscope images of centromeres of PD cells. Quantification of changes in centromeres areas is shown in Figure 5A. **B:** Diplotene cells were stained for H3K9me3. Areas of chromocenters of diplotene cells were calculated using NIS elements software and expressed as a fraction of the mean of all experimental groups in arbitrary units (a.u). Each data point represents the total area of one cell stained with H3K9me3 antibody, mean±SE of about 80 cells in each group, one-way Anova, Tukey's multiple comparisons. **C**. Histone lysine modifications were measured using a histone extraction kit ab113476 and multiplex colorimetric assay kit ab185910, 25 ng protein/well, extracts were prepared from vehicle and oxamate incubated cells, from two differentexperiments. **D**. Cells in diakinesis (evident by typical SCP3 pattern of staining) were imaged. White circles mark paired centromeres. Representative images are shown. **E.** Number of centromere pairs in diakinesis. 30-50 cells were evaluated for number of pairs of centromeres per nucleus. Mean±SE; Tukey's multiple comparisons test.

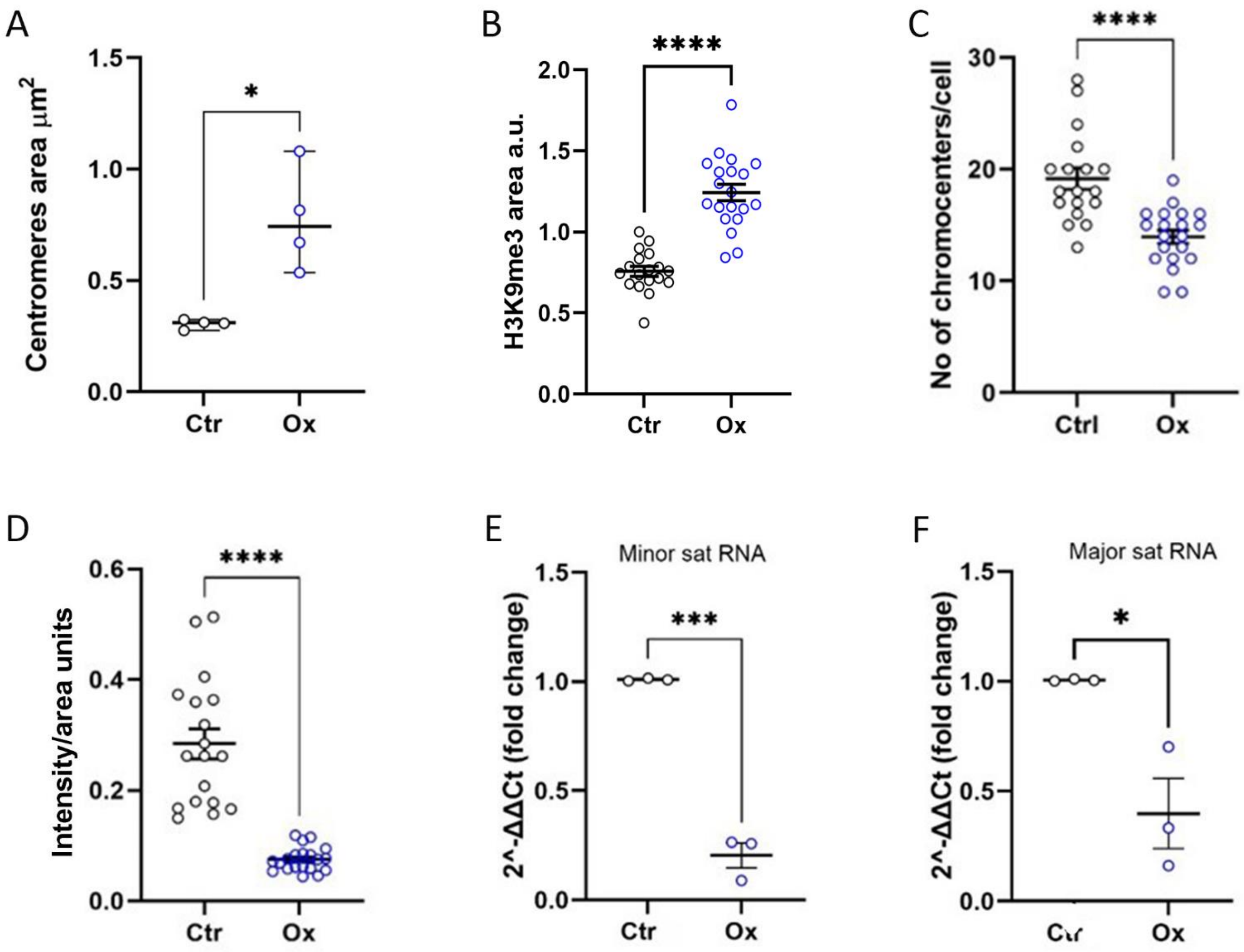


**Figure S4: Oxamate treatment of mice recapitulates effects on centromeres and chromocenters noted in the *in vitro* setting.** For **A-D** mice were injected once with oxamate 1.3g/Kg or vehicle; for **E-F** similar injections were done for two successive days. All mice were killed 24 hrs after oxamate injection and PDD cells were isolated. **A.** PDD nuclear spreads were stained with CREST antibodies. Centromere size of diplotene cells was quantified with NIS elements analyser software. Each data point represents the mean of >200 ROIs from one experiment. Median with range of 4 experiments, Mann Whitney. **B**. Cells were stained with H3K9me3 antibody. Area of chromocenters of diplotene cells was calculated using NIS elements software and expressed as a fraction of the mean of both experimental groups in arbitrary units (a.u). Each data point represents the total area of one cell , mean±SE, of about 20 cells in each group. **C**. Numbers of chromocenters were counted for each diplotene cell using data from **B**. **D**. Total intensity of red fluorescent stain of H3K9me3 / area unit /cell. **B-D**: Mean ±SE-t-test. **E-F.** Minor (**E**) and major (**F**) satellite RNA was quantified with qPCR. Mean±SE of 3 independent experiments; t-test.

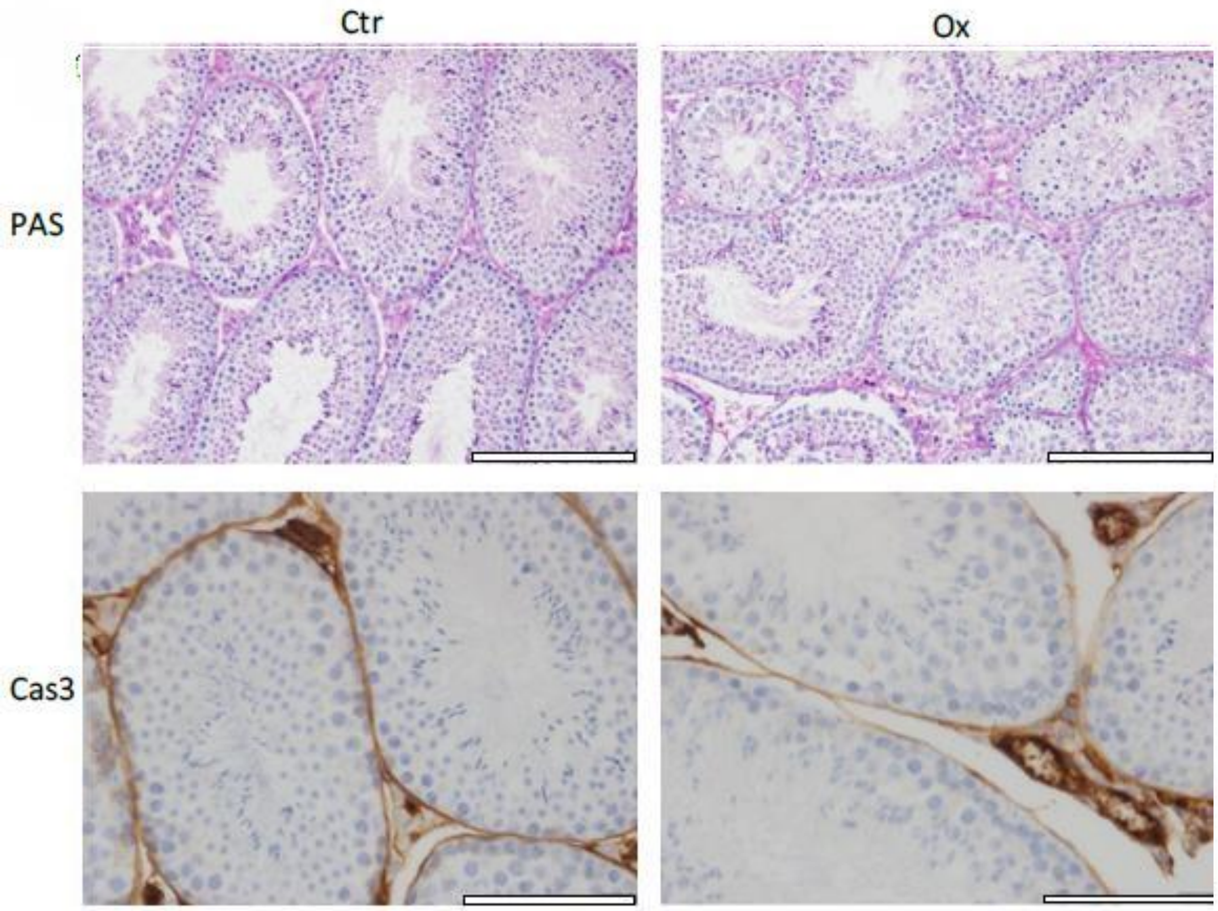


**Figure S5: Lack of histological signs of toxicity after 7 days of treatment with oxamate:** Mice were treated with daily oxamate injections 1.3 g/kg or vehicle for 7 days (n=6). Paraffin sections were stained with PAS for morphological evaluation or with cleaved caspase 3 for identification of apoptotic cells. Representative images one out of 6 mice.

| | Diameter µm | Radius µm | Surface Area $\mu m^2$ | Volume $\mu m^3$ |
|---|---|---|---|---|
| ***Eclipse*** | | | | |
| Spg | 12.4 | 6.2 | 486 | 1008 |
| LZ | 9.9 | 5.0 | 308 | 509 |
| PD | 16.9 | 8.4 | 895 | 2518 |
| RS | 11.3 | 5.6 | 398 | 747 |
| Microscope | | | | |
| Spg | 13.2 | 6.6 | 544 | 1195 |
| LZ | 10.6 | 5.3 | 351 | 618 |
| PD | 18.0 | 9.0 | 1022 | 3073 |
| RS | 12.9 | 6.5 | 526 | 1136 |
| Average of ***Eclipse*** and Microscope | | | | |
| Spg | 12.8 | 6.4 | 515 | 1099 |
| LZ | 10.2 | 5.1 | 329 | 562 |
| PD | 17.5 | 8.7 | 957 | 2786 |
| RS | 12.1 | 6.1 | 460 | 928 |

**Table S1: Cell size parameters of the different cell populations.** Cell volumes were measured using two independent methods: eclipse and diameter measurement under the microscope, with high level of agreement between the two methods. The volume ratio between the respective cells was calculated based on the average of the two methods.
